# Retracing and rewriting the evolutionary trajectories of mammalian developmental enhancers

**DOI:** 10.64898/2026.04.20.719714

**Authors:** Tony Li, Jean-Benoît Lalanne, Emma A.N. Kajiwara, Shruti Jain, Xiaoyi Li, Tiffany V. Do, Beth K. Martin, Samuel G. Regalado, Riza M. Daza, Jay Shendure

## Abstract

*Cis*-regulatory elements (CREs) such as enhancers play a central role in orchestrating mammalian development, yet how they have gained, lost, maintained or changed function over the course of mammalian evolution remains poorly understood. To address this gap, we densely mapped the functional evolution of five mouse developmental enhancers by testing orthologous sequences from 480 extant and ancestrally reconstructed mammalian genomes (Zoonomia^1^, Cactus^2^) with massively parallel reporter assays (MPRAs). This phylogenetic dissection revealed diverse modes of evolution, from lineage-restricted activity to deep functional conservation despite extensive sequence divergence. To pinpoint causal changes, we developed a model-driven reconstitution strategy that uses deep learning-based predictions of chromatin accessibility to re-introduce a succession of mutations into ancestral orthologs; this revealed critical transcription factor binding site (TFBS) changes and pervasive context-dependent epistasis, including instances where mutational effects were strongly contingent on the order of their introduction. When we extended this strategy to tune the activity of extant orthologs, we found that ablation of enhancer function required as few as one to seven mutations, whereas enhancement was constrained by element-specific activity ceilings—a striking asymmetry in the predictability of model-guided enhancer editing. Together, these results shed light on how the plasticity of mammalian enhancers intersects with their evolution, and advance a framework for reprogramming the activity of endogenous CREs at nucleotide resolution.

## INTRODUCTION

Mammalian CREs orchestrate the spatiotemporal and cell type-specific gene expression programs that underlie the transformation of a zygote into a complex organism. While the core *trans*-regulatory networks specifying cell identities are deeply conserved, CREs evolve rapidly and are thought to be the primary drivers of species-specific phenotypic divergence^3–9^. Even single nucleotide changes in CREs can reshape developmental programs and underlie evolutionary innovations^3,4,10–13^. Conversely, many CREs maintain regulatory function despite extensive sequence divergence^14–16^.

Although genome-wide comparative studies have revealed broad patterns of mammalian enhancer evolution^7,17–21^, the specific sequence changes that shaped the evolutionary trajectories of individual CREs are largely unexplored^11,22^. Seminal studies have shown that many conserved human enhancers exhibit similar expression patterns to their evolutionary orthologs (via transgenic mice)^20,23^, and that numerous cell type-specific enhancers have been gained and lost throughout mammalian evolution (via biochemical profiling of extant mammalian cell lines)^7,24^. Yet these approaches fall short of a fine-grained, nucleotide-resolution understanding of how specific sequence changes shape the gain, loss, or maintenance of enhancer function across the mammalian phylogeny. More recently, deep learning models have emerged as powerful tools for interpreting the regulatory grammar of cell type-specific CREs^25,26^, but their potential to illuminate enhancer evolution remains largely untapped^27,28^.

Although the resurrection and functional characterization of ancient proteins has yielded fascinating insights into protein evolution^29,30^, only a few studies have extended this paradigm to non-coding regulatory elements^31,32^. Importantly, a similar asymmetry between sequence and functional evolution operates in the regulatory realm: while enhancer sequences change rapidly, the trans-regulatory networks that interact with them — the transcription factors and chromatin modifiers that shape genomic output — are deeply conserved^5^ and cell type-specific rather than species-specific, so much so that human chromosomes transplanted to mice exhibit chromatin landscapes that are practically indistinguishable in orthologous cell types^6^.

The stark mismatch in pace of *cis* vs. *trans* evolution creates an opportunity for investigating how CREs evolve — specifically, by transplanting many extant and ancestral orthologous CREs into a common model system representing the *trans*-regulatory environment of a cell type of interest. Historically, a barrier to this paradigm has been insufficient power to accurately infer ancestral enhancer sequences. However, Christmas *et al.* recently reported the Zoonomia resource, a set of high-quality reference genomes for 241 extant mammalian species^1^. Through the combination of Zoonomia and Cactus^2^, a reference-free multiple genome alignment program, it is now possible to access not only extant orthologs of a given CRE, but also computationally reconstructed sequences of putative ancestral orthologs at each internal node of the mammalian phylogeny. When combined with modern experimental and computational tools for scalably quantifying regulatory activity, we are newly in a position to systematically investigate the fine-scale evolution of CREs throughout the mammalian phylogeny (**Fig. 1a**).

**Figure 1.**
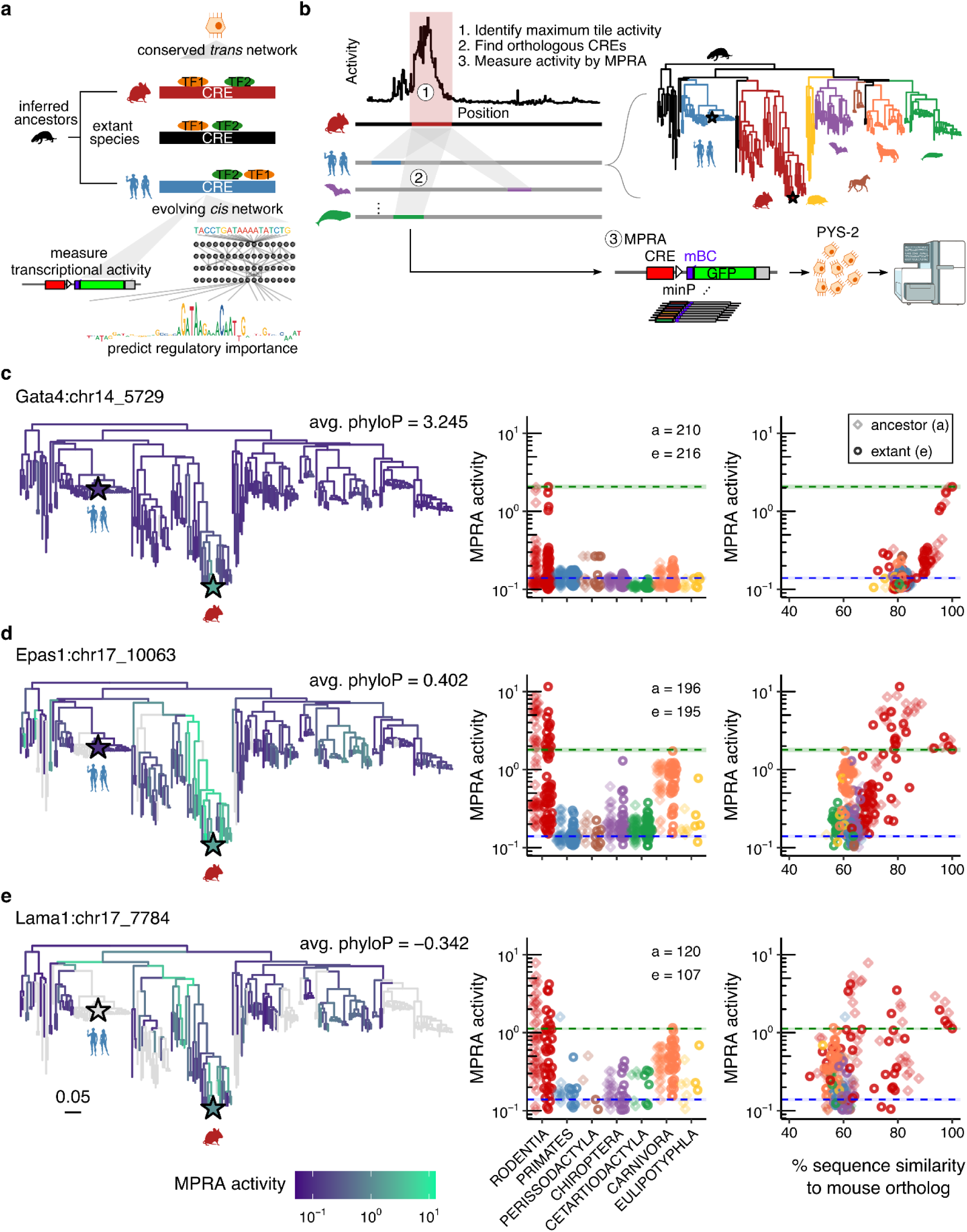
Functional profiling of ancestral and extant orthologs of five mouse parietal endoderm enhancers. **a)** Schematic of the strategy for investigating CRE evolution by leveraging the deep conservation of cell type-specific *trans-*regulatory networks. Ancestral and extant orthologs of a cell type-specific CRE are functionally profiled in extant cells of that type, followed by deep learning-based modeling to nominate and interpret functional sequence changes. **b)** Schematic of experimental workflow. For each CRE, the maximum activity tile was identified by sub-tiling MPRA^38^, and ancestral and extant orthologs were retrieved from Cactus alignments and ancestral reconstructions of 241 mammalian genomes. Orthologs were synthesized, cloned upstream of a minimal promoter driving a barcoded GFP reporter, and functionally characterized by bulk MPRA in mouse PYS-2 cells, which model the *trans*-regulatory environment of the mammalian parietal endoderm. Filled and open stars indicate the positions of mouse and human, respectively, on the evolutionary tree. **c-e)** Left: Mammalian phylogenetic tree with branches colored by the MPRA-measured activity of ancestral or extant orthologs of the *Gata4* **(c)**, *Epas1* **(d)**, and *Lama1* **(e)** proximal CREs in PYS-2 cells. MPRA activities represent normalized RNA/DNA ratios. Grey branches denote species for which the ortholog was either absent from the Cactus alignments or not recovered in the assay. Filled and open stars denote the activities of mouse and human orthologs, respectively. The mean phyloP conservation score across the mouse ortholog sequence is shown. A scale bar indicates branch length in substitutions per site. Middle: MPRA activity broken out by taxonomic order, with the numbers of ancestral (a) and extant (e) orthologs listed. Right: MPRA activity as a function of percent sequence similarity to the mouse ortholog. In both middle and right panels, green and blue dotted lines indicate the activity of the mouse ortholog and minP background control, respectively; lighter and darker hues denote ancestral and extant orthologs, respectively.

Here we set out to test this paradigm by leveraging a mouse cell line to model the *trans-*regulatory environment of the parietal endoderm, a major subtype of the extraembryonic endoderm that plays a key role in early embryogenesis and for which a well-characterized cell line surrogate is available. Focusing on five developmental enhancers whose extant mouse orthologs exhibit high activity and specificity in this cell type in embryoid bodies, we profiled 1,871 extant or ancestrally reconstructed orthologs from 480 mammalian genomes using massively parallel reporter assays (MPRAs), which enable simultaneous functional measurement of thousands of sequences in a single experiment^33^. Our results reveal a diversity of evolutionary trajectories, including a *Gata4*-proximal element that exhibits lineage-restricted activity despite deep sequence conservation, and *Epas1-* and *Lama1-*proximal elements that maintain activity across distant clades despite extensive sequence divergence. Finally, to pinpoint causal changes underlying gain, loss or change of activity, we developed a reconstitution framework that leverages deep learning models of chromatin accessibility — which capture cell type-specific regulatory grammar at nucleotide resolution — to nominate the order in which mutations are introduced into inferred ancestral sequences, with functional impact then validated by MPRAs. We then applied this framework to tune the activity of extant enhancers, and found that function can be substantially enhanced or entirely ablated with a handful of model-nominated mutations. Taken together, our results illuminate how mammalian enhancers gain, lose, and maintain function over evolutionary timescales, and establish a paradigm in which synthetic genomics and sequence-based models together enable the experimental traversal of cell type-specific regulatory sequence space.

## RESULTS

### Selection of five parietal endoderm-specific CREs and validation of a surrogate cell system

We recently developed the single-cell expression quantitative reporter (scQer) assay and applied it to mouse embryoid bodies (mEBs) to identify 13 cell type-specific developmental CREs, 8 of which were exclusively active in parietal endoderm^34^. As quantitative profiling of large numbers of CREs with scQer remains challenging due to the throughput and cost constraints of single-cell profiling, we sought to establish a simpler system in which these autonomously active, parietal endoderm-specific CREs, each approximately 1 kb in length, could be tested with a conventional bulk MPRA.

For this purpose, we selected the PYS-2 cell line, which derives from the yolk sac of a mouse teratocarcinoma and has been used to model the mammalian extraembryonic endoderm^35,36^. Comparison of genome-wide chromatin accessibility between PYS-2 (via bulk ATAC-seq) and various mEB germ layers (via pseudobulk ATAC-seq) revealed strong correlation with parietal endoderm cells (Spearman’s *ρ* = 0.64) but weak correlation with other lineages (*ρ* = 0.20-0.26; **Fig. S1a**). This specificity was even stronger for differentially accessible peaks (*ρ* = 0.71 for parietal endoderm vs. *ρ* < 0 for other lineages). Comparison of the PYS-2 chromatin accessibility profile to *in vivo* data from gastrulating mouse embryos^37^ again showed the strongest correlation with parietal endoderm (*ρ* = 0.64 vs. *ρ* = 0.11-0.37 for other cell types; **Fig. S1b**). Finally, the eight scQer-validated CREs were robustly accessible in PYS-2, while *Sox2*-proximal, pluripotent stem cell-specific CREs were inaccessible (**Fig. S1c**).

Together, these analyses establish PYS-2 cells as a reliable surrogate for the chromatin accessibility landscape of the mouse parietal endoderm lineage. Bulk RNA-seq of PYS-2 cells was also most correlated with pseudobulk parietal endoderm among mEB cell types (*ρ* = 0.86 vs. *ρ* = 0.80 for other cell types; **Fig. S1d**), and only parietal endoderm exhibited strong expression of the core endodermal transcription factors Foxa2, Klf4, Gata4/6, and Sox17^38^.

To pinpoint where regulatory activity is localized within each of these ∼1 kb CREs, we performed a sub-tiling MPRA experiment, systematically tiling all eight CRE sequences into 270-bp windows with 5-bp steps^38^. For five of the eight CREs, the maximal subsequence activity coincided with the peak summit of our mouse parietal endoderm scATAC-seq data. These five CREs are located near genes with established roles in parietal endoderm differentiation, including early-acting transcription factors *(Gata4) a*nd late-stage effectors (*Lama1*, *Sparc*)^39,40^, among others. As each of these CREs (*Gata4*:chr14_5729, *Epas1*:chr17_10063, *Lama1*:chr17_7784, *Sparc*:chr11_7211, and *Bend5*:chr4_8201) exhibited strong activity in PYS-2 cells and were accessible in both PYS-2 and mEB-derived parietal endoderm cells, we prioritized their extant and ancestral sequences for comparative and evolutionary analysis across mammals. In a companion manuscript^38^, these same five enhancers are deeply dissected via a complementary strategy of multi-scale dissection, compaction and derivatization (Lalanne, Li *et al.* <u>bioRxiv</u> 2026).

### Phylogenetic distributions of activity for 1,871 extant and ancestral enhancer orthologs

We systematically profiled the phylogenetic activity distributions of extant and ancestral orthologs of these five mouse CREs by leveraging Zoonomia genomes^17^ and their associated Cactus alignments^2^. For each CRE, we used *halLiftOver*^2^ to map a 300-bp window centered on the maximally active tile to 240 extant and 239 ancestrally reconstructed mammalian genomes, generating a catalog of 1,871 CRE orthologs spanning the full diversity of mammalian lineages for functional testing in PYS-2 cells (**Fig. 1b**). Phylogenetic coverage varied widely, from 55.3% for Lama1 CREs to 99.6% for Gata4 CREs (**Table S1**), likely reflecting differential loss of orthologs through deletion^41^.

To functionally characterize this library, we employed bulk episomal 5’ MPRAs in PYS-2 cells, in which the relative activities of thousands of CREs positioned upstream of a minimal promoter (minP) are quantified by counts of CRE-associated barcodes in RNA, normalized to counts of those same barcodes in DNA^22,42^ (**Fig. S2a**). Following synthesis, cloning, and CRE-barcode association, the library contained 1,867 of the 1,871 nominated CRE orthologs (99.8%), with 195 ± 140 barcodes per CRE (n = 290,862 barcodes altogether). Library recovery was high and consistent across replicates (DNA: mean 99.4% barcodes recovered; mean *ρ* = 0.92 for log-transformed counts between biological replicates; **Fig. S2b**), and the assay spanned nearly three orders of magnitude of activity between negative (minP only, 𝜇 = 0.14, 𝝈 = 0.006) and positive (EEF1A1p, 𝜇 = 74.9, 𝝈 = 1.0) controls, with an internal IGVF promoter standard series^43,44^ confirming high reproducibility and dynamic range (**Fig. S2c**). As a further validation that PYS-2 cells recapitulate the relevant *trans*-regulatory environment, six of the eight scQer-validated parietal endoderm-specific CREs robustly drove expression in PYS-2 cells. The most active of these, the *Epas1-*proximal CRE (𝜇 = 1.03; 𝝈 = 0.02), exceeded background levels by ∼7-fold, while no activity was detected for the two pluripotent stem cell-specific CREs (𝜇 < 0.1; **Fig. S2d**).

We then mapped the resulting activities across all orthologs onto the mammalian phylogeny (**Figs. 1c-e**; **Figs. S3a-c**). CRE orthologs corresponding to extant and ancestral rodents generally retained similar activity to the extant mouse CRE ortholog, consistent with their low sequence divergence. However, we observed diverse patterns for larger evolutionary distances. *Gata4* CRE orthologs rapidly declined in activity as a function of sequence divergence, despite being the most conserved of the five elements (**Figs. 1c**, **S3a**; median sequence divergence of orthologs retaining >25% of mouse activity: 18.7%). In contrast, many *Epas1* and *Lama1* CRE orthologs retained robust activity despite substantially greater divergence (median divergences: 33% and 37%, respectively; **Figs. 1d-e**, **S3a**). This was largely driven by coherent pockets of activity in distant sub-clades — the *Epas1* and *Lama1* CREs were robustly active in Carnivora and Chiroptera lineages, respectively (**Fig. 1d-e**). When broken out by taxonomic order, correlations between sequence identity and functional activity ranged from *ρ* = −0.76 to 0.79 (**Fig. S3d**). Strikingly, the *Gata4* CRE — the most conserved of the five elements, with detectable orthology even in the chicken genome (mean phyloP = 3.25; **Fig. S4a**) — rapidly loses activity beyond rodents, while the poorly conserved *Epas1* and *Lama1* CREs (mean phyloP = 0.40 and-0.34, respectively) retain robust activity across several distant clades. Together, these patterns reveal a clear decoupling of sequence and functional conservation that is both element-and lineage-specific, and cannot be captured by simple conservation-based models of regulatory function.

To assess whether this decoupling extends to base-pair resolution, we performed saturation mutagenesis MPRAs on all five CREs^38^ and compared mutational effects to phyloP scores. Mutational sensitivity showed no overall correlation with conservation across elements (*ρ* < 0.02; **Fig. S4b**). and while functional TFBSs were generally more conserved than non-functional positions, this enrichment was confined to the *Gata4* and *Epas1* CREs (**Fig. S4c**). These findings reinforce the conclusion that sequence constraint is an unreliable guide to regulatory function, and motivate a more direct approach to mapping how specific sequence changes shape enhancer activity across the mammalian phylogeny.

### Mutational basis of a *Mus* genus-specific gain-of-function in the *Gata4* CRE

To investigate the specific sequence changes underlying the functional trajectories observed above, we examined the relationship between stepwise changes in CRE activity and the emergence or affinity maturation of endoderm-specific TFBSs along inferred evolutionary paths (**Fig. 2a**). We focused on the *Gata4* and *Epas1* CREs, which were the best-represented of our five elements in mammalian Cactus alignments (∼200 ancestral and ∼200 extant orthologs each; **Fig. 1c-d**), and examine each in turn below.

**Figure 2.**
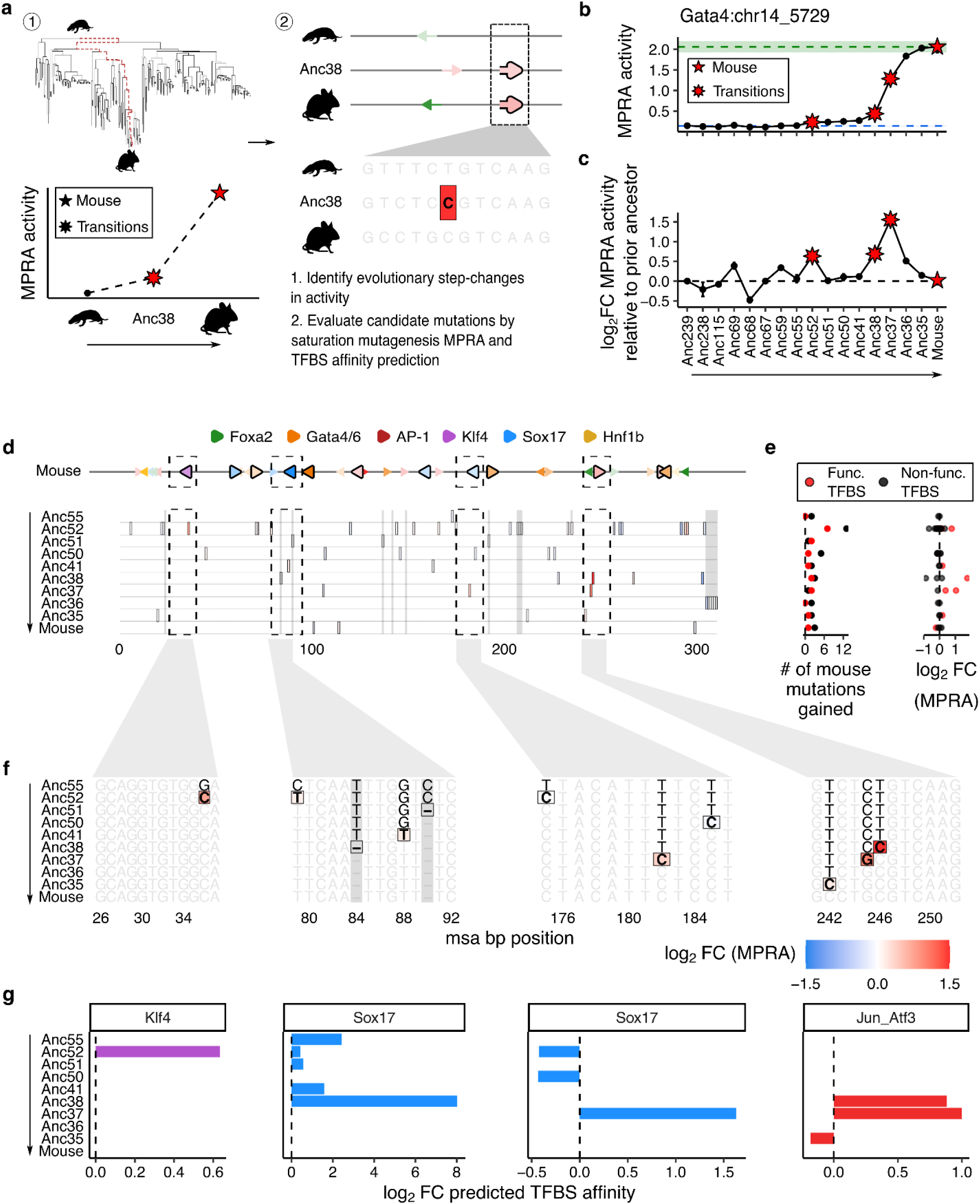
Mutational basis of a Mus-restricted gain-of-function in the Gata4 CRE. **a)** Schematic of strategy. We identified step changes in MPRA activity of the *Gata4* CRE along the evolutionary path from the inferred common mammalian ancestor to the present-day mouse, then leveraged saturation mutagenesis MPRA data and TFBS affinity predictions to identify which derived mutations may have driven each step change. Filled and asterisk red stars denote the extant mouse ortholog and key ancestral transitions, respectively. **b-c)** Absolute MPRA activity **(b)** and log₂ fold-changes in MPRA activity relative to the immediately prior ancestor **(c)** along the evolutionary trajectory of the *Gata4* CRE leading to *M. musculus*; star symbols as in panel **a**. Error bars indicate standard error across three experimental replicates. **d)** Multiple sequence alignment (msa) of *Gata4* CRE sequences along this evolutionary trajectory. Predicted TFBS for the mouse ortholog are shown at the top; those functional by saturation mutagenesis MPRA are indicated by larger triangles with black outlines. In the msa, black-bordered tiles correspond to derived mutations relative to the immediately prior ancestor, colored by log_2_ fold-changes in saturation mutagenesis MPRA activity (blue = decreased activity, red = increased activity; scale shown in panel **f**). Gray columns mark derived indels not assessed by saturation mutagenesis MPRA. Black dashed boxes highlight four functionally supported TFBSs further detailed in panels **f** and **g**. **e)** Number (left) and magnitude of log_2_ fold-changes (right) of derived mouse mutations appearing at each evolutionary step, partitioned by whether mutations lie within (red) or outside (black) a functional mouse TFBS. **f)** Nucleotide resolution views of the msa from panel **d**, highlighting mutations within four functional TFBS that are candidate drivers for the three largest step changes. Bolded, boxed nucleotides indicate the first appearance of a derived mutation, colored by log_2_ fold-change in saturation mutagenesis MPRA activity (color scale at bottom; blue = decreased activity, red = increased activity); unbolded, unboxed nucleotides indicate the ancestral state. Gray columns mark derived indels not assessed by saturation mutagenesis MPRA. **g)** Log_2_ fold-change in predicted TFBS affinity at each evolutionary step for the four functional TFBSs highlighted in panel **f**. Negative values indicate predicted loss of affinity relative to the immediately prior ancestor; positive values indicate predicted gain.

Outside of the rodent lineage, ancestral and extant *Gata4* CRE orthologs do not drive expression in PYS-2 cells (**Fig. 1c**), despite the deep sequence conservation of this element across mammals (**Fig. S4a**) and a high density of conserved, functionally relevant TFBS^38^ (**Fig. S5a**). This suggests that the *Gata4* CRE retains an ancestral regulatory function in some other cell type context, and that its activity in parietal endoderm is a derived feature specific to the rodent lineage — and as we show below, largely to the *Mus* genus.

What mutations underlie the 14.1-fold gain-of-function for the *Gata4* CRE in PYS-2 cells accruing between the inferred common mammalian ancestor (Anc239) vs. extant *Mus musculus* (**Fig. 2b**)? We focused on the three largest step changes: the 1.6-fold, 1.6-fold and 3.0-fold gains in activity at Anc52, Anc38, and Anc37 relative to their immediately prior ancestral nodes, which collectively account for half of the 14.1-fold gain (**Fig. 2c**). Notably, although Anc38 corresponds to the inferred common ancestor of mouse and rat, further gains did not accrue along the path to the extant rat ortholog, whose activity is 8.8-fold lower than that of the extant mouse ortholog and 1.9-fold lower than Anc38 (**Fig. S5b-c**), consistent with a gain-of-function that is largely specific to the *Mus* genus (**Fig. S5d**).

For each of these three step changes, we examined mutations acquired along the path from Anc55 to the extant mouse *Gata4* CRE. The first step change, between Anc55 and Anc52, is coincident with 20 ancestral-to-derived changes — defined as mutations newly present in Anc52 that are also retained in the extant mouse ortholog — (Anc52 row of **Fig. 2d**-**f**). To evaluate their potential functional effects, we leveraged the saturation mutagenesis MPRA of the mouse *Gata4* CRE, which correlated strongly with predicted TFBS affinities^38^, enabling the mapping of twelve high-confidence functional binding sites in this sequence. Of the 20 mutations distinguishing Anc55 from Anc52, seven fall within these predicted extant TFBSs, but nearly all have little impact on enhancer activity (MPRA mean log₂FC = 0.11). The exception was a G→C substitution within a predicted extant Klf4 TFBS (msa position 36) which leads to a 1.7-fold gain in activity by MPRA (Anc52 row of **Fig. 2d-f**) and coincides with a 1.6-fold gain in predicted Klf4 binding affinity (Klf4-focused subpanel of **Fig. 2g**). A single mutation may thus account for the full 1.6-fold Anc55→Anc52 step change (with the caveat that the saturation mutagenesis MPRA tests each mutation on the background of the extant mouse sequence rather than the ancestral sequence, which limits the precision of this inference).

The two remaining step changes were coincident with fewer mutations, making their interpretation more tractable. The Anc41→Anc38 transition coincided with only 5 derived mutations, and Anc38→Anc37 with only 3. Four of these eight mutations occurred within three functional extant TFBSs — two Sox17 and one AP-1 site (**Fig. 2d-f**). As with the first step change, both saturation mutagenesis data and predicted binding affinities pinpointed specific mutations as likely drivers of the corresponding gains in activity. For the Anc41→Anc38 step change: a deletion of G at position 84 was associated with a 260-fold gain in predicted Sox17 affinity, and a T→C substitution at position 246 with a 1.8-fold gain in predicted AP-1 affinity and a 3.4-fold gain in MPRA activity. For the Anc38→Anc37 step change: a T→C substitution at position 182 was associated with a 3.1-fold gain in predicted Sox17 affinity and a 1.3-fold gain in MPRA activity, and a C→G substitution at position 245 with a 2.0-fold gain in predicted AP-1 affinity and a 2.1-fold gain in MPRA activity (**Fig. 2d-f**; Sox17 and AP-1-focused subpanels of **Fig. 2g**).

The extensive conservation of the *Gata4* CRE, beyond the positions highlighted by this ancestral retracing, indicates that this element retains a deeply conserved regulatory function in some non-parietal endoderm context across Mammalia. Against this backdrop, the dramatic gain in parietal endoderm activity in a restricted clade — largely driven by a handful of mutations affecting Klf4, Sox17, and AP-1 binding sites — may represent a striking example of enhancer repurposing. These mutations appear to have accrued gradually over tens of millions of years, either creating new TFBSs or enhancing the affinity of previously weak sites, collectively enabling the *Gata4* CRE to acquire parietal endoderm regulatory activity within *Mus*.

### Fragility and plasticity of cis-regulatory modules shape *Epas1* CRE activity across mammals

While the *Gata4* CRE’s activity is largely confined to the *Mus* genus, the *Epas1* CRE exhibited robust activity more broadly — throughout Rodentia as well as throughout several non-rodent clades (**Fig. 1d**). To identify sequence features underlying this complex phylogenetic distribution, we examined saturation mutagenesis data for the mouse *Epas1* CRE^38^, which highlighted two TFBS triplets as essential for its function: (i) a heterotypic module of Foxa2-Sox17-Gata4/6 sites, and (ii) a homotypic module of AP-1 sites (**Fig. 3a**). We therefore sought to trace the emergence and turnover of motifs within these modules throughout mammalian evolution, using the same ancestral-path framework applied to the *Gata4* CRE above.

**Figure 3.**
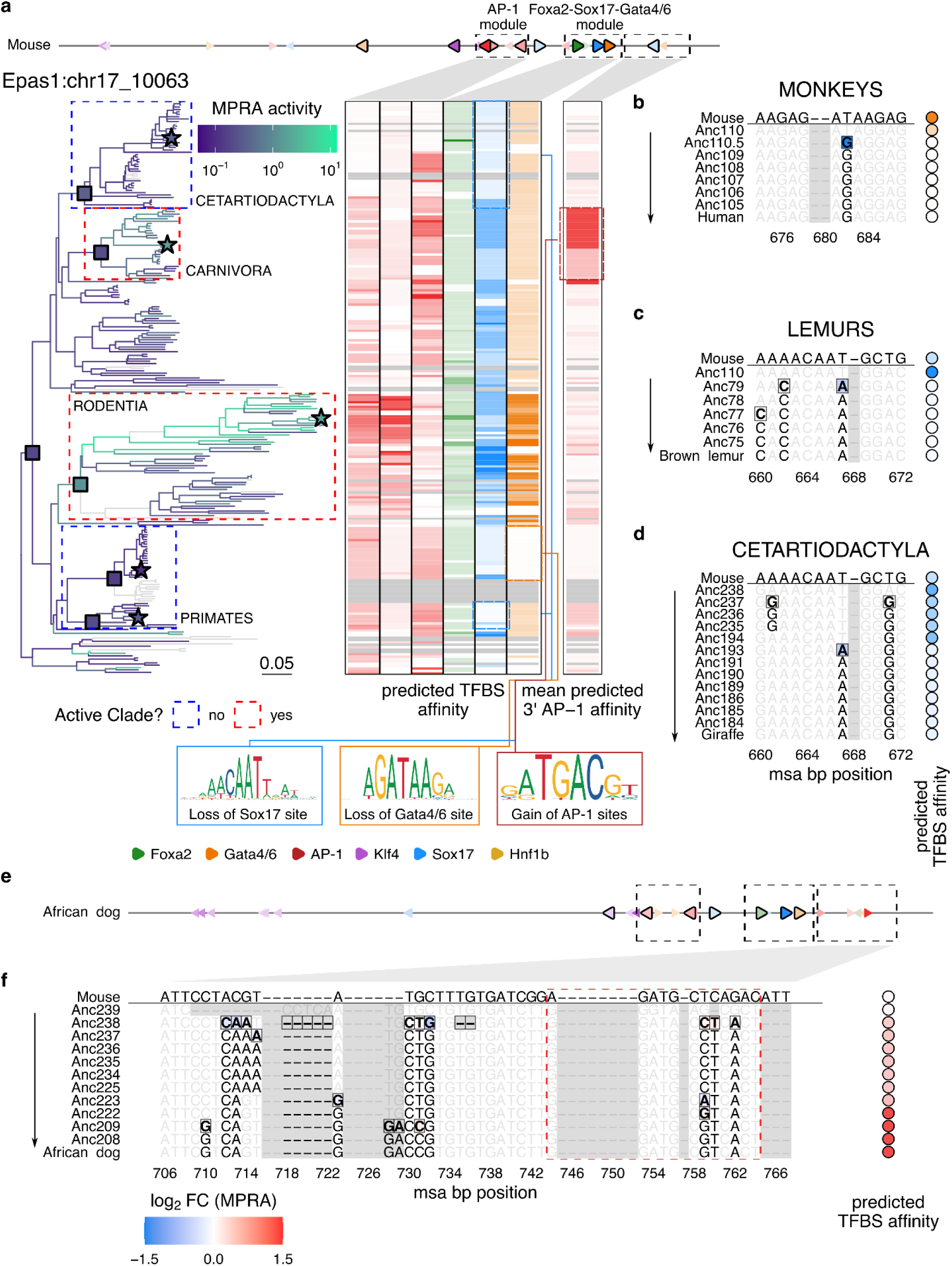
Fragility and plasticity of *cis*-regulatory modules shape *Epas1* CRE activity across mammals. **a)** Heatmap of predicted TFBS affinities for all extant mammalian *Epas1* CRE orthologs, highlighting the homotypic AP-1 module, the heterotypic Foxa2-Sox17-Gata4/6 module, and a Carnivora-specific AP-1 site. For each TFBS, predicted affinities are normalized across orthologs and color-coded by relative value. Predicted TFBS positions for the extant mouse ortholog are shown at top, with black dashed boxes denoting the modules visualized in the heatmap. At left, the mammalian phylogeny is colored by MPRA activity (as in **Fig. 1d**); filled squares mark key ancestral clade nodes, including Anc238 (Boreoeutherian), Anc223 (Carnivora), Anc194 (Cetartiodactyla), Anc110 (Primates), and Anc68 (Rodentia). Dashed boxes highlight clades that gain (red) or lose (blue) activity; stars mark the extant species highlighted in panels **b-f**. Sequence logos at the bottom illustrate representative motif changes associated with loss of the Sox17 site, loss of the Gata4/6 site, and gain of AP-1 sites, in the indicated clades. **b-d)** Nucleotide-resolution MSAs for individual TFBSs within the conserved triplet module, tracing evolutionary paths from selected ancestral to extant orthologs: **b,** Gata4/6 TFBS along the path from Anc110 → *H. sapiens* (human); **c**, Sox17 TFBS along the path from Anc110 → *E. fulvus* (brown lemur); **d**, Sox17 TFBS along the path from Anc238 → *G. tippelskirchi* (giraffe). In each panel, bolded, boxed nucleotides denote the first appearance of substitutions relative to Anc110 or Anc238, colored by log₂ fold-change in saturation-mutagenesis MPRA activity^38^ (color scale at bottom left of panel **f**; measured in mouse background. Unbolded, unboxed nucleotides indicate the Anc110 or Anc238 state; gray columns mark derived indels not assayed by MPRA. Right side bars indicate TFBS identity (colors as in the TF legend in panel **a**) and predicted TFBS affinity (hue intensity), normalized across sequences shown. The extant mouse ortholog sequence is shown above each MSA for reference. **e)** Predicted TFBS positions for African wild dog (*L. pictus*) ortholog. Black dashed boxes correspond to the modules highlighted in panel **a**. **f)** Nucleotide-resolution MSA of the 3′ end of the *Epas1* CRE, tracing the path from Anc239 → *L. pictus*, highlighting emergence of a Carnivora-specific AP-1 site. Presentation follows panels **b-d**. Red dashed box highlights 3’ AP-1 site exhibiting gains in predicted affinity. In the right side bar, hues correspond to predicted AP-1 affinity, normalized across the set of sequences shown, colored by log₂ fold-change in saturation-mutagenesis MPRA activity^38^ (color scale at bottom left; measured in mouse background).

Tracing the evolutionary emergence of the Foxa2-Sox17-Gata4/6 module along the lineage leading to extant mouse, we identified four mutations first appearing in the inferred Boreoeutherian ancestor, Anc238 that collectively established the full triplet motif (**Fig. S6a-e**). When introduced individually into the extant mouse ortholog, substitutions from the ancestral to derived states at these positions were associated with 18-fold to 338-fold gains in predicted TFBS affinity, and 3-fold to 12-fold gains in MPRA activity. Once established in Anc238, this module was stably maintained throughout the ancestral lineage leading to extant mouse, and exhibited elevated sequence conservation relative to other functional TFBSs within the *Epas1* CRE (mean phyloP of 1.5 vs. 0.4; **Fig. S6f**).

To assess the conservation of this regulatory module at higher resolution, we evaluated its integrity across all *Epas1* CRE orthologs with respect to both predicted TFBS affinity and MPRA-measured activity. While several clades retained intact versions of each TFBS comprising the heterotypic triplet, others showed reduced affinity or complete loss of the Sox17 or Gata4/6 binding sites, or both (**Fig. 3a**). Within the primate order alone, we identified two independent losses of enhancer function — one in old world monkeys and another in lemurs — each associated with mutations predicted to disrupt the Foxa2-Sox17-Gata4/6 module. In the monkey subclade, a T→G substitution in the Gata4/6 motif (msa position 682), present in old world monkeys including humans (**Fig. 3b**, Anc110.5 row), was predicted to reduce Gata4/6 binding affinity by 41-fold and, when introduced into the mouse *Epas1* CRE, reduced enhancer activity by 5.0-fold (**Fig. 3b**). In the lemur clade, two substitutions arising between Anc110 and Anc79 — A→C (pos 662) and T→A (pos 667) — were together predicted to weaken Sox17 binding affinity by 29-fold (**Fig. 3c**). Notably, the same T→A mutation at position 667 was inferred to have arisen independently in Anc193, a reconstructed ancestor within the Cetartiodactyla order; in the giraffe *Epas1* ortholog, this substitution combined with a T→G (pos 671) substitution in Anc237 was predicted to reduce Sox17 binding affinity by 6.4-fold (**Fig. 3d**).

In these three independent examples, loss of just one motif within the Foxa2-Sox17-Gata4/6 heterotypic triplet was sufficient to completely abrogate enhancer activity. All *Epas1* CRE orthologs were inactive in the clades downstream of these mutations, whereas activity was retained or emergent in adjacent clades where the module remained intact (**Figs. 1d**, **3a**). Although these patterns establish the triplet module as necessary for *Epas1* CRE function, this module in isolation is insufficient to drive measurable activity in PYS-2 cells^38^. Consistent with this, the inferred Boreoeutherian ancestor, Anc238, despite having acquired the full heterotypic triplet, was similarly inactive (**Figs. S7a-b**).

What other sequences are necessary for the *Epas1* CRE to be active in parietal endoderm? We observed that the homotypic AP-1 trio, the second module essential for *Epas1* CRE function in mouse, appeared specifically within the rodent lineage (**Figs. 3a**, **S6a**). This cluster was established through a series of insertions and substitutions (msa positions 181–194) beginning at Anc69, the inferred ancestor of the Glires grandorder (rodents and lagomorphs; **Fig. S7c-d**). Of particular note, a T→A mutation (pos 182) in Anc59 created the central AP-1 motif and was associated with a marked increase in enhancer activity. These mutations were retained in the extant mouse *Epas1* CRE, and, in combination with the Foxa2–Sox17–Gata4/6 module, are sufficient to reconstitute its full activity^38^.

Although both the Foxa2-Sox17-Gata4/6 and rodent-specific AP-1 triplets are required for robust *Epas1* CRE activity in Rodentia, orthologs from other parts of the mammalian phylogeny — most strikingly a subclade of Carnivora that includes African wild dogs — were also consistently active (**Figs. 1d**, **S7e-f**). Do the coherent patterns of activity beyond Rodentia exhibit the same dependencies? In this Carnivora subclade, the Foxa2-Sox17-Gata4/6 module and two of three AP-1 motifs were intact, but the central AP-1 motif critical for activity in rodents was absent^38^ (**Figs. 3e**, **S7g**). Upon closer examination, we identified a distinct AP-1 site located on the other side of the heterotypic triplet whose emergence was specific to this Carnivora subclade. Specifically, the combination of C→T mutation (pos 760) appearing in Anc238 and a T→G mutation (pos 759) in Anc222 — the inferred ancestor of the high-activity Carnivora subclade — was predicted to increase AP-1 binding affinity by 23-fold (**Fig. 3a**,**f**).

In summary, the *Epas1* CRE illustrates a striking combination of fragility and flexibility. Our data suggest that the Foxa2-Sox17-Gata4/6 heterotypic triplet module arose in the inferred Boreoeutherian ancestor and is preserved in most but not all of its extant descendants. The module appears necessary for enhancer function, as disruption of any one of its three motifs through as few as one or two mutations abrogates activity potential in three independent clades—Old World monkeys, lemurs, and giraffes. Yet in lineages that retained the module, such as rodents and carnivores, strong activity in parietal endoderm further depended on the emergence of AP-1 motifs. In rodents, a cluster of three adjacent AP-1 motifs arose through successive mutations within the Glires lineage; in carnivores, a distinct AP-1 site was gained at the opposite end of the element, achieving a functionally similar outcome. Altogether, these diverse patterns of gain and loss at just this single CRE illustrate how dynamic the evolution of mammalian enhancers can be, with subtleties that are unlikely to be inferred from sequence conservation patterns alone.

### Synthetic reconstitution of gain-of-function trajectories reveals epistasis and TFBS affinity tuning

Having identified candidate TFBS mutations underlying the major gains in parietal endoderm activity of these CREs, we next asked whether these changes are sufficient to functionally reconstitute enhancer activity from an inactive ancestral sequence, or whether additional sequence context is required. To nominate candidate mutational trajectories for experimental testing, we evaluated several sequence-based deep learning models (**Supplementary Note 1; Extended Data Figs. 1-3**) and selected ChromBPNet^45,46^, which is trained to predict chromatin accessibility rather than MPRA activity, yet exhibited the strongest correspondence with MPRA measurements while also providing base-resolved estimates of the effect of individual nucleotide substitutions.

We focused this analysis on the *Gata4*, *Epas1*, and *Lama1* CREs. We began by enumerating sequence differences between the inactive ancestral ortholog (Anc239) and the active extant mouse ortholog; pairwise alignments revealed 53, 94, and 111 substitutions and indels, corresponding to sequence differences of 13%, 22%, and 26% for the *Gata4*, *Epas1*, and *Lama1* CREs, respectively. We then iteratively added these derived mutations into the ancestral ortholog, using ChromBPNet at each step to select the mutation predicted to most strongly increase chromatin accessibility (**Fig. 4a**). This yielded a stepwise trajectory from Anc239 → mouse for each CRE (navy lines in **Fig. S8a**). In parallel, we generated 10 control trajectories per CRE in which the same mutations were introduced in a random order (light gray lines in **Fig. S8a**).

**Figure 4.**
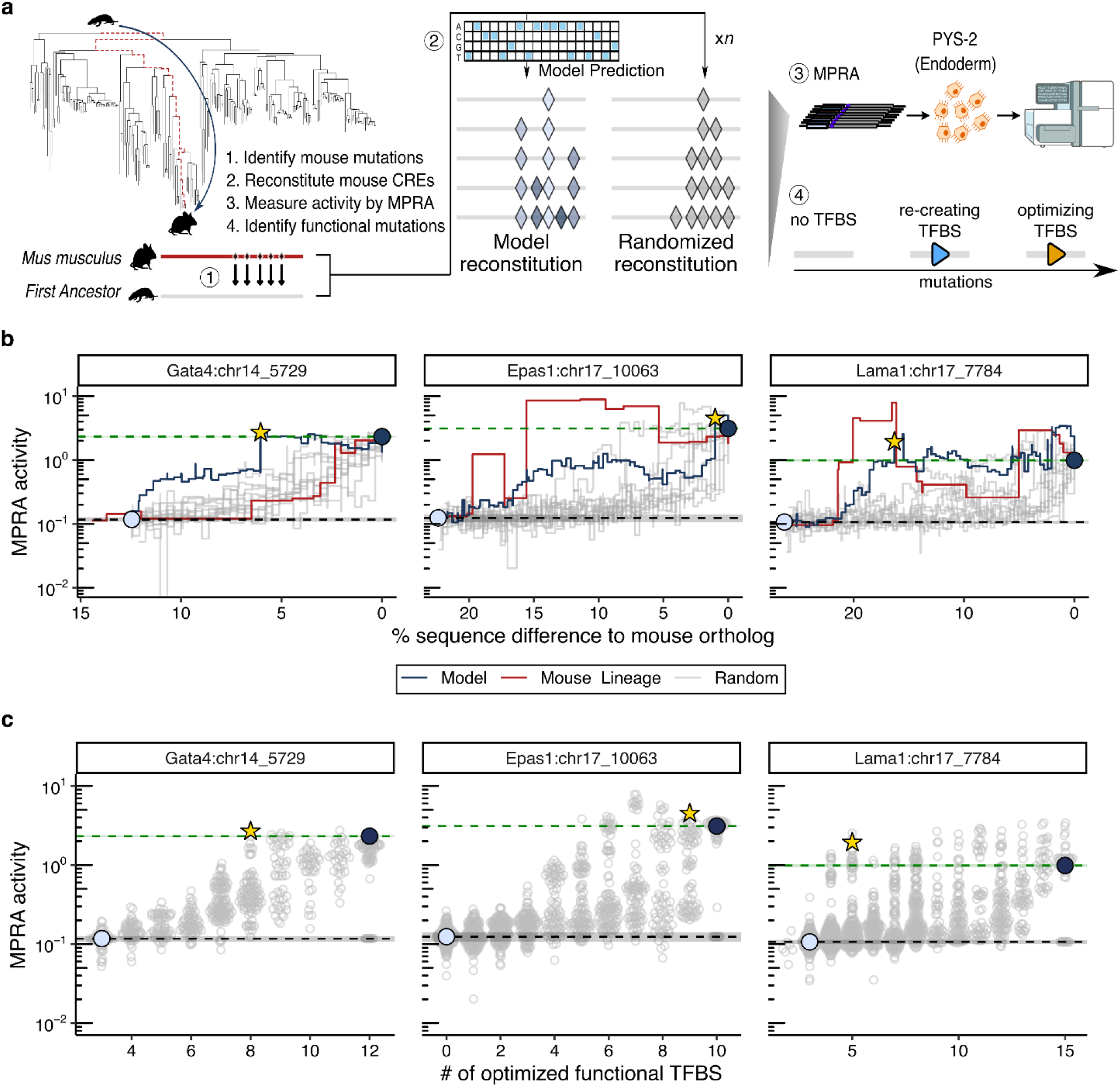
Synthetic reconstitution of enhancer gain-of-function trajectories reveals epistasis and TFBS affinity tuning. **a)** Schematic of the model-guided reconstitution strategy. Starting from the inferred common mammalian ancestral ortholog (Anc239), which is inactive in PYS-2 cells, mouse-derived substitutions and indels are sequentially introduced either in a ChromBPNet-optimized order or in a random order to generate alternative mutational trajectories toward the extant mouse CRE. **b)** MPRA-measured activity (y-axis) as a function of sequence divergence from the extant mouse ortholog (x-axis) for reconstitution trajectories of the *Gata4*, *Epas1*, and *Lama1* CREs. Model-optimized, random-order, and phylogeny-inferred evolutionary trajectories are shown in navy, light gray, and red, respectively. Circles indicate the Anc239 starting point (light blue) and extant mouse endpoint (navy) for all trajectories. Stars denote the earliest intermediate along the model-optimized trajectory whose activity equals or exceeds that of the endogenous mouse CRE. Green and black dotted lines indicate the activity of the extant mouse ortholog and the inferred common mammalian ancestral ortholog, respectively. **c)** MPRA activity (y-axis) as a function of the number of fully reconstituted functional TFBSs (x-axis), defined as sites whose predicted binding affinity equals or exceeds that of the corresponding TFBS in the endogenous mouse CRE. Dotted lines, circles, and stars are as in panel **b**.

These *in silico* trajectories predicted that reintroducing a small subset of mouse-derived mutations into inactive Anc239 orthologs is sufficient to recover the predicted accessibility levels of extant mouse orthologs (9, 19, and 10 mutations for the *Gata4*, *Epas1*, and *Lama1* CREs, respectively); the model also predicted intermediates whose accessibility greatly exceeded that of the extant mouse (**Fig. S8a**). In contrast, randomized trajectories required reintroduction of nearly all mutations to achieve comparable accessibility. The predicted accessibility trajectories of phylogeny-informed mutational orders fell within the distribution of random orders for the *Gata4* and *Lama1* CREs, but more closely matched the model-optimized trajectory for the *Epas1* CRE (red lines in **Fig. S8a**).

Next, we experimentally evaluated both the model-optimized and random mutational trajectories by functionally assaying each intermediate via bulk MPRA in PYS-2 cells. Across the *Gata4*, *Epas1*, and *Lama1* CREs, we synthesized, cloned, and tested 2,788 of 2,959 (94%) intermediate sequences (**Fig. 4b**). Although MPRA-measured activity broadly agreed with predicted accessibility (*ρ* = 0.78–0.82; **Fig. S8b**), experimental trajectories differed substantially from model predictions, likely reflecting both the limitations of model performance and of accessibility as a proxy for transcriptional output^47^ (**Fig. S8b** vs. **Fig. 4b**). Nonetheless, model-optimized trajectories outperformed nearly all random trajectories — random orders typically required reintroduction of nearly all mouse-derived mutations to restore full activity, although a small number of random *Epas1* trajectories achieved functional recovery as rapidly as, or even faster than, the model-optimized path (light gray vs. navy lines in **Fig. 4b**).

Notably, ancestral intermediates along the inferred evolutionary trajectories for the *Epas1* and *Lama1* CREs reached or exceeded mouse-level activity earlier than synthetic trajectories at equivalent sequence divergence, indicating that high activity is achievable on sequence backgrounds other than that of the extant mouse ortholog (red lines in **Fig. 4b**). In contrast, *Gata4* CRE ancestral intermediates underperformed the model-optimized trajectory, suggesting that the *Gata4* CRE’s relatively recent gain of parietal endoderm activity in *Mus* was dependent not only on specific TFBS mutations, but also the broader sequence context in which those mutations occurred (**Fig. 1c**)

To what extent do stepwise functional gains along these trajectories correspond to the creation of new TFBSs, the quantitative optimization of existing TFBSs, or changes to background sequence? We quantified the fractional change in MPRA activity associated with each mutation introduced along each reconstitution trajectory and mapped these effects relative to functional TFBSs in the mouse reference sequence. For the *Gata4* and *Epas1* CREs, mutations overlapping extant TFBSs tended to have larger effect sizes (**Fig. S9a**), and for all three CREs, TFBS-overlapping mutations accounted for a disproportionate share of functional recovery — though this share was more variable for the *Epas1* CRE than the *Gata4* and *Lama1* CREs (**Figs. S9b-c**; observed vs. expected: 82 ± 14% vs. 42% for *Gata4*; 67 ± 32% vs. 41% for *Epas1*; 67 ± 11% vs. 55% for *Lama1*). Upon partitioning mutations into those that first reconstituted a functional TFBS, but possibly at a lower affinity than in the endogenous endpoint, versus those that optimized the affinity of an existing TFBS, we found that activity reconstitution was mainly driven by TFBS-optimizing mutations (**Fig. S9d**). Consistent with this, the optimization of as few as 8 of 12 (*Gata4* CRE), 5 of 12 (*Epas1* CRE) or 4 of 15 (*Lama1* CRE) functional TFBS to affinities matching or exceeding their wildtype affinities was sufficient to restore endogenous levels of activities in at least one model-optimized or randomized trajectory (**Fig. 4c**).

Does the order in which mutations are introduced modulate their contribution to functional recovery? For each mutation appearing across both model-optimized and random trajectories, we quantified how functional impact varied by order of introduction, and compared this order-dependent variability to that observed across technical replicates of the same trajectory. For all three CREs, order-dependent variability in mutational effect sizes was 2-3-fold greater than experimental noise, indicating that epistatic interactions among evolutionarily derived mutations substantially influenced functional trajectories — and by extension, may have influenced the historical paths by which these enhancers gained activity during mammalian evolution..

In summary, synthetic reconstitution offers a means of contextualizing mutational trajectories, whether evolutionarily plausible, model-optimized or random, with respect to their optimality and dependencies. Across elements, functional recovery was driven disproportionately by mutations that tune the affinity of functional TFBSs, and the effect of any given mutation depended strongly on the sequence context in which it was introduced. Together, these results help define the mutational paths through regulatory sequence space that lead to gains in enhancer activity, and establish reconstitution trajectories as a practical benchmark for sequence-based models — one that requires models to correctly predict not just endpoints, but the ordering, context dependence, and diminishing returns that govern bona fide mutational paths.

### Model-guided tuning of endogenous activity reveals asymmetry between enhancement and ablation

Having shown that deep learning can guide the stepwise reconstitution of enhancer activity from inactive ancestral sequences, we next asked whether the same framework could be applied to tune the activity of endogenous enhancers. While deep learning models have been leveraged to propose gain-of-function edits in inactive sequences^48^, their ability to guide the modulation of endogenous CRE activity upwards (enhancement) or downwards (ablation) has not been systematically evaluated.

To this end, we implemented a gradient-based strategy using ChromBPNet to iteratively nominate *de novo* substitutions predicted to either increase or decrease parietal endoderm chromatin accessibility across our five endogenous CREs (**Fig. 5a**). For each CRE, the model predicted monotonic accessibility trajectories under each objective, in some cases predicting up to 100-fold changes relative to the predicted accessibility of the extant mouse sequence (**Fig. S10a**). A null model trained on pluripotent chromatin accessibility did not recapitulate these predictions for the same mutated sequences, consistent with ChromBPNet having learned regulatory features specific to the parietal endoderm program rather than general chromatin accessibility rules (**Fig. S10a**).

**Figure 5.**
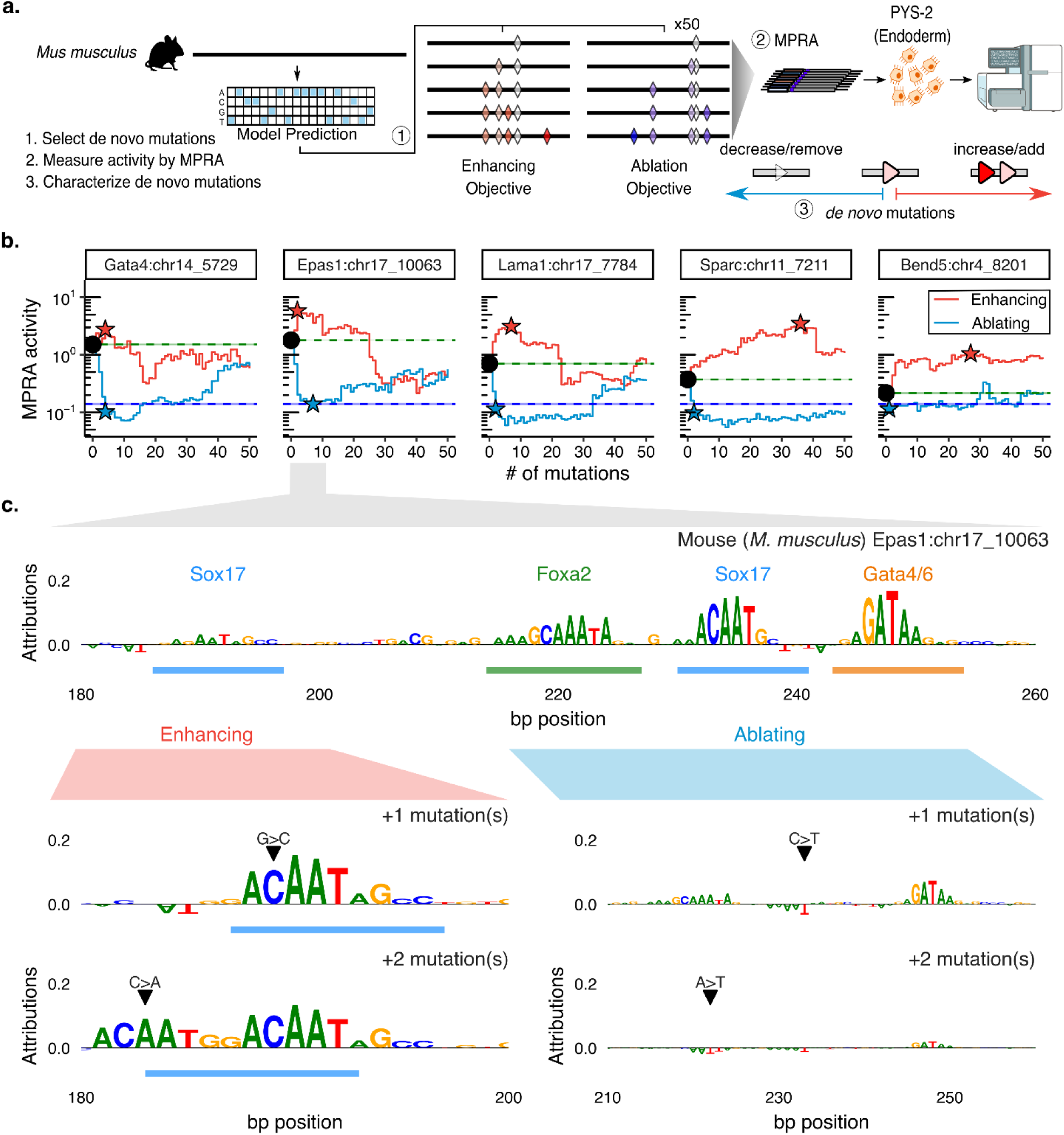
Model-guided enhancement and ablation trajectories of endogenous parietal endoderm CRE activity. **a)** Schematic of the model-guided tuning strategy. For each mouse CRE, *de novo* single-nucleotide substitutions are iteratively nominated by ChromBPNet to either maximally increase (enhancement objective) or maximally decrease (ablation objective) predicted chromatin accessibility, for up to 50 iterative steps. **b)** MPRA-measured activity of intermediate sequences along the model-guided trajectories for each CRE, shown separately for enhancement (red) and ablation (blue) objectives. Points denote MPRA activity at each step; black circles indicate the starting endogenous sequence and stars indicate the intermediate achieving the greatest enhancement or ablation of activity relative to the endogenous sequence. Horizontal dotted lines (with shading) denote the activity of the endogenous mouse CRE (green) and the minP background control (blue). **c)** ChromBPNet per-nucleotide contribution scores, reflecting the predicted influence of each base on chromatin accessibility, shown across positions 180–260 of the mouse *Epas1* CRE. Colored bars indicate functional TFBSs defined by saturation mutagenesis. Bottom panels show contribution scores after introducing the first two mutations selected under the enhancement objective (left; positions 180–200) or the ablation objective (right; positions 210–260), illustrating how the model’s predicted regulatory grammar shifts upon mutation. Black arrows mark the positions and identities of the introduced substitutions.

We then synthesized and measured transcriptional output for the first 50 steps of each model-nominated enhancement or ablation trajectory (**Fig. 5a**). Functional testing of these trajectories by MPRA revealed a strong asymmetry between ablation and enhancement. For all five CREs, activity dropped to negative-control levels with just 1 to 7 mutations under the ablation objective, corresponding to 1.9- to 14.8-fold reductions from endogenous activity (**Fig. 5b**).

Under the enhancement objective, however, the achievable upregulation appeared to be a function of baseline activity. CREs with lower baseline activity (*Sparc*, *Bend5*) mirrored the monotonic enhancements predicted by the model (*ρ* > 0.4), achieving peak activity after dozens of model-nominated mutations had been introduced (36 and 27, respectively) that collectively achieved major gains in activity (9.3- and 4.7-fold, respectively) (**Figs. 5b**, **S10b**). In contrast, CREs with robust baseline activity (*Gata4*, *Epas1*, *Lama1*) exhibited rapid functional saturation, peaking after a few model-nominated mutations (4, 2, and 7, respectively) and more modest gains in activity (1.8-, 3.2- and 4.4-fold, respectively) (**Fig. 5b**). For these elements, agreement between predicted accessibility and MPRA activity degraded early (*ρ* < −0.5; **Fig. S10b**), suggesting rapid divergence from the sequence contexts represented in the model’s training data^49,50^.

Given the rapid departure from the model’s training distribution after a limited number of mutations, we focused subsequent analyses on the first ten mutations within each enhancement or ablation trajectory. Across all five CREs, these early model-nominated mutations were distributed across both functional TFBS (mean 44%) and non-TFBS (mean 56%) positions (**Fig. S11a**), suggesting that the model targets a mix of binding site and background sequence features. Notably, under the ablation objective, mutations disrupting functional TFBSs produced significantly larger decreases in activity than non-TFBS mutations (up to 5-fold; p < 0.01; **Fig. S11b**), consistent with TFBS disruption being the primary driver of the rapid ablation observed above.

As a concrete illustration of model behavior, we examined the initial mutations selected for enhancement and ablation of the *Epas1* CRE, whose regulatory modules are well-characterized from the preceding sections. Under the enhancement objective, two substitutions within a functional Sox17 TFBS (G→C at position 190 and C→A at position 184) were associated with a 5.8-fold increase in predicted Sox17 binding affinity and together increased MPRA activity by 3.3-fold (**Fig. 5c**, left), consistent with previous observations that tuning the affinity of key TFBS can robustly boost enhancer output^51^. Under the ablation objective, the model instead targeted the Foxa2–Sox17–Gata4/6 module, where disrupting the Sox17 and Foxa2 binding sites with substitutions (C→T at position 234 and A→T at position 223) weakened predicted binding affinities by 7.1- and 18.2-fold, respectively, and together reduced MPRA activity by 8.9-fold (**Fig. 5c**, right).

Finally, to assess whether the effects of successive mutations combined additively or cooperatively, we used single-mutation effect sizes from our saturation mutagenesis data^38^ to predict the combined effects of multiple mutations under additive and multiplicative models, and compared these predictions to the observed MPRA trajectories (**Fig. S11c**). Across all five CREs, an additive model explained a substantial fraction of observed activity changes (<u>ρ</u> = 0.61–0.88; **Fig. S11d**). However, for the *Gata4* and *Epas1* CREs, a multiplicative model provided a superior fit (<u>ρ</u> = 0.75–0.78), suggesting that even short mutational trajectories can produce cooperative interactions whose combined effects exceed simple additive expectations.

In summary, model-guided ablation of endogenous enhancer activity was achieved with as few as 1 to 7 mutations, primarily at TFBSs, whereas enhancement was constrained by element-specific activity ceilings that the model failed to anticipate. Together, these results reveal a pronounced asymmetry in the reliability of current sequence-to-function models for tuning endogenous enhancers: ablation trajectories are short and predictable, whereas enhancement trajectories saturate and then diverge from model expectations. This divergence becomes most evident as engineered sequences move farther from the manifold of naturally evolved regulatory sequences, exposing practical limits of model-guided enhancer optimization, and suggesting a need for models trained on broader and more diverse sequence contexts.

## DISCUSSION

Enhancers outnumber genes by several orders of magnitude and play a central role in shaping mammalian development, evolution and phenotypic variation^3^. Yet how the stepwise accumulation of noncoding mutations intersects with a slowly evolving *trans-*acting regulatory milieu^52^ to drive the gain, loss, or maintenance of cell type-specific activities over evolutionary timescales remains underexplored, usually investigated through the study of extant orthologs^14,15,53–57^, rather than ancestral intermediates^31,32^. Here, taking a page from the protein ancestral reconstruction playbook^29,30^, we provide what is, to our knowledge, the most fine-grained investigation to date of how mutational trajectories shape the evolution of mammalian enhancers. Profiling both extant and ancestral orthologs across the mammalian phylogeny allows us to move beyond endpoint comparisons and trace the relationship between sequence evolution and functional evolution.

We intentionally pursued the deeper dissection of a small number of endogenous developmental enhancers, rather than a broader but shallower survey, resulting in element-specific insights into how the stepwise accumulation of mutations underlies functional change. The *Gata4* CRE illustrates how deep sequence conservation can coexist with lineage-specific regulatory innovation. Despite conservation across and beyond mammals, its parietal endoderm activity is restricted to the *Mus* lineage, arising from a handful of mutations affecting Klf4, Sox17, and AP-1 binding sites that accrued gradually over tens of millions of years. This raises the intriguing possibility that CREs conserved for their role in one cell type context may commonly harbor latent regulatory potential for other contexts, which may be selectively co-opted in particular lineages^54^.

In contrast, the *Epas1* CRE demonstrates how enhancer function can be robust to extensive sequence remodeling, provided that a core regulatory module is preserved. Disruption of this module — through as few as one or two mutations — leads to a collapse of regulatory potential in three independent clades, whereas its retention enables coherent pockets of activity across distant mammalian lineages through the independent emergence of distinct AP-1 architectures: a homotypic AP-1 triplet in rodents, and a distinct 3’ AP-1 site in carnivores, each arriving at a functionally similar outcome by a different mutational route. Together, the *Gata4* and *Epas1* histories illustrate how the same ancestral regulatory scaffold can be repurposed through alternative motif solutions or dismantled by just a few mutations — dynamics that may be more widespread across the repertoire of developmental mammalian enhancers^7,10,13–16^.

The synthetic reconstitution trajectories provide a complementary perspective on the functional landscape available to endogenous enhancers; specifically, what stepwise mutational routes are ‘accessible’ to them. By ordering naturally occurring mutations into ChromBPNet-optimized paths from inactive ancestral sequences to active extant orthologs, we show that equivalent gains are achievable with far fewer mutations than actually occurred. Similar to evolutionary trajectories, these ‘shortcuts’ are disproportionately reliant on the affinity maturation of existing TFBSs rather than the creation of new ones. However, they also highlight a strong dependence on the order of introduction, suggesting pervasive epistasis that extends beyond key TFBS. A similar conclusion is reached in our companion manuscript, in which a complementary strategy of multi-scale dissection, compaction and derivatization revealed pervasive background sequence effects^38^.

What happens when deep learning-based regulatory models are asked to predict or design not only endpoints^49,50^, but also paths from point A to point B? By densely characterizing potential mutational trajectories whose starting point is an extant or ancestral endogenous enhancer, we provide challenging scenarios for predictive frameworks. The divergence between predicted and measured activity after a modest number of mutations is notably asymmetric: models reliably predict ablation trajectories, where disrupting a handful of TFBSs is sufficient and predictable, but fail to anticipate element-specific activity ceilings. This asymmetry likely reflects the fact that ablation operates near the training distribution whereas enhancement quickly ventures into sequence regimes that are increasingly distant from the naturally evolved enhancer sequences on which models are trained. Rather than failures, these discrepancies are informative boundaries that expose where current representations of regulatory grammar remain untested

Three limitations warrant emphasis. First, our functional profiling was conducted in PYS-2 cells, which we validated as a reliable surrogate for the parietal endoderm trans-regulatory environment. However, embryoid bodies — from which the original scQer data were derived^34^ — do not recapitulate all developmental contexts, and our results suggest that at least some of the CREs studied here are active in additional cell type contexts. The *Gata4* CRE is the clearest example: its deep sequence conservation, including at positions not highlighted by saturation mutagenesis in PYS-2 cells, implies a conserved function elsewhere that our experimental system cannot capture. This is arguably a strength as much as a limitation, as functional profiling across phylogenies provides a means to dissect how enhancers evolve when they serve multiple regulatory roles, as many enhancers likely do. Looking forward, the challenge of cell type context dependence may be addressable through complementary approaches that attempt to model regulatory activity across all developmental contexts^58^.

A second limitation is that the ancestral sequences used here are computational reconstructions^2^ and therefore subject to inference uncertainty. While our analyses are likely robust to modest sequence variation, inaccuracies arising from limitations in genome quality or taxon sampling could affect inferred ancestral states and the activities we attribute to them, a caveat that may be remedied as the corpus of extant mammalian genomes grows further, enabling more accurate reconstructions.

Finally, we are limited by our use of chromatin accessibility as a surrogate for transcriptional activity in our deep learning model-guided synthetic trajectories. While accessibility has been shown to be a strong predictor for regulatory activity^48,59,60^, these are not equivalent phenomena, and a model trained on ATAC-seq signal is unlikely to fully capture the determinants of cell type-specific transcriptional output. Future iterations of this framework may benefit from models that are either trained directly or fine-tuned on activity readouts^61–63^.

The coupling of synthetic genomics, ancestral inference, and sequence-based modeling enables the explicit traversal of mutational paths through regulatory sequence space — revealing which changes are sufficient, necessary, or contingent on prior context, and allowing us to reorder naturally occurring mutations to test how alternative paths affect regulatory output. As both DNA synthesis technologies^64,65^ and sequence-based regulatory models^66^ continue to advance, we anticipate that the field will progress from enhancers to loci to genomes^64,67^, positioning this paradigm not merely as a complement to comparative genomics, but also as a more general means of reconstructing ancestral programs of gene and developmental regulation.

## AUTHOR INFORMATION

## Acknowledgments

We thank the members of the Shendure Lab, particularly the ‘gene regulation’ subgroup including F. Abadie, D. Calderon, W. Chen, and T. McDiarmid, for extensive and helpful discussions, as well as X. Li, W. Yang, C. Suiter, C. Kubo, D. Lee, C. Qiu, Q. Yu, and H. Liao. We also thank the Ahituv lab, Kircher lab and R. Waterston for helpful discussions. This work was supported by the National Human Genome Research Institute (UM1HG011966 to J.S.), the Brotman Baty Institute for Precision Medicine and the Seattle Hub for Synthetic Biology, a collaboration between the Allen Institute, Chan Zuckerberg Initiative and University of Washington (award number CZIF2023-008738 to J.S.). J.-B.L. was supported by a Damon Runyon Cancer Research Foundation fellowship (DRG-2435-21) and a Next-Generation Scientist award from the Cancer Research Society (grant no. 1155581). J.S. is an Investigator of the Howard Hughes Medical Institute.

## Author Contributions

Conceptualization, T.L., J.B.L., and J.S.; Investigation, T.L. and J.B.L.; Data Curation, T.L., J.B.L., E.A.N.K., T.V.D., S.J., and X.L.; Formal Analysis, T.L., J.B.L., and S.J.; Visualization, T.L. and J.B.L.; Resources, B.K.M., S.G.R., R.M.D., and J.S.; Manuscript writing, T.L., J.B.L., and J.S.; Funding Acquisition, J.S.; Supervision, J.S.

## Data and Code Availability

Raw and processed sequencing data generated in this study have been deposited and are freely available, including MPRA data (IGVF portal https://data.igvf.org/, accession numbers IGVFDS9730EWQE and IGVFDS1754MDNZ) and RNA/ATAC data (IGVF portal, accession numbers IGVFDS8171UAYI and IGVFDS6861BLHS), custom sequencing amplicon data for CRE-barcode associations (GEO, accession number GSE328309). Single-cell ATAC-seq data used to train the chromatin accessibility model were previously published^34^ (GEO, accession number GSE217683). Single cell RNA-seq from the previous dataset is also used (GEO, accession GSE217686). Single-cell ATAC-seq data from *in vivo* gastrulating mouse embryos were previously published^37^ (GEO, accession number, GSE205117). Code to reproduce results and generate figures is available at https://github.com/shendurelab/mammalian-cre-evolution-and-design.

## AI Disclosure Statement

We disclose that language editing, proofreading and coding were supported by AI-based tools; these were not used for conceptual development or primary manuscript writing. The authors take full responsibility for the contents of this manuscript.

## METHODS

### Cell culture

PYS-2 cells (CRL-2745, ATCC) were grown in Dulbecco’s modified Eagle medium (DMEM; Thermo Fisher, cat. no. 11995065), supplemented with 10% FBS (Fisher Scientific, Cytiva HyClone fetal bovine serum, cat. no. SH3039603) and 1× penicillin/streptomycin (Thermo Fisher, cat. no. 15140122). Cells were kept at 37 °C and 5% CO2, and passaged every 2 days.

### Bulk RNA sequencing and processing

Bulk RNA sequencing of PYS-2 cells was performed using a commercially available service (Plasmidsaurus). To meet shipping requirements, cells were split into four technical replicates, each consisting of approximately 100,000 cells resuspended in 100 µL of Zymo DNA/RNA Shield. For downstream analyses, FASTQ files from each replicate were aligned to the mouse reference genome (GRCm39/mm39) using *bowtie2*, with alignments processed using *samtools* under default parameters. Gene-level count matrices were generated using the *featureCounts* function from the *Subread* package with default settings. To facilitate comparison with previously generated single-cell RNA-seq data, count matrices from biological replicates in the scRNA-seq dataset were first stratified by cell type and then averaged within each cell type prior to downstream analyses.

### Bulk ATAC sequencing and processing

Nuclei were isolated from PYS-2 cells and formaldehyde-fixed as previously described^68^. Briefly, fixation was performed by adding 37% formaldehyde to a final concentration of 1% for 10 min at room temperature, and the reaction was quenched with 2.5 M glycine. Fixed nuclei were resuspended in freezing buffer (50 mM Tris pH 8.0, 25% glycerol, 5 mM MgOAc₂, 0.1 mM EDTA, 5 mM DTT, and 1x protease inhibitors) at ∼1M nuclei per aliquot and stored at −80°C. Three frozen vials derived from the same plate were processed independently as technical replicates.

Frozen nuclei were thawed on ice, pelleted at 500 x g for 5 min at 4°C, and repermeabilized in 100 µL Omni lysis buffer on ice for 3 min. The reaction was then neutralized by addition of 500 µL RSB-Tween^69^. Nuclei were counted on a hemocytometer with trypan blue, and 50,000 nuclei were used per tagmentation reaction. Tagmentation was performed in Nextera TD buffer with 1X DPBS, 0.01% Digitonin, 0.1% Tween-20 and 2.5 μl Nextera v2 enzyme (Illumina) per reaction, and incubated at 55°C for 30 min. To stop tagmentation, reactions were supplemented with EDTA (20 mM final concentration) and incubated at 37°C for 15 min. Nuclei were then pelleted at 500 x g for 5 min at 4°C and resuspended in 20 µL Qiagen elution buffer (EB). Crosslinks were reversed by adding Proteinase K (0.83 mg/mL final, from 20 mg/mL stock) and SDS (0.04% final, from 1% stock), followed by incubation at 65°C for 16 hours.

Reactions were purified using the Zymo DNA Clean & Concentrator kit with 2x binding buffer, and eluted in 22 µL Qiagen EB. Tagmented DNA was amplified with NEBNext HF 2x PCR Mastermix, BSA and indexed P5/P7 oligos. Amplification was monitored in real time and stopped at 8 cycles to preserve library complexity and avoid over-amplification bias. Amplified libraries were size-selected by double-sided SPRI cleanup (0.5x/1.0x) to remove large fragments and primer dimers, respectively. Final libraries were sequenced on Illumina NextSeq 2000 platform using a P2 100-cycle kit with 59/10/10/59 read configuration.

For the bulk ATAC-seq analysis of the PYS-2 dataset, we used the ENCODE pipeline^19^ to generate irreproducible discovery rate (IDR) peaks and corresponding BAM files. Bigwig pileups were generated by merging replicate BAM files using *samtools*, followed by bedtools^70^ *genomecov* and *wigToBigWig* from kentUtils^71^.

### Single cell chromatin accessibility data preprocessing

Biological replicates from published scATAC-seq datasets^34^ (GSE217690) were processed using *subset-bam_linux* to separate the BAM files by cell types, which were identified using cell barcodes. After splitting the BAM files by cell type, we merged the replicate BAM files, and the resulting files were sorted and indexed using *samtools*^72^ v1.21. Peaks for each cell-specific bam file were identified using macs2^73^ *callpeak* function (options: --shift-100 --extsize 200-p 0.01-g 1.87e9). The peaks were filtered by removing those overlapping the ENCODE mouse blacklisted regions, which were extended by 1057 bp on both sides. Additionally, we excluded any peaks overlapping with the tested elements from our previous study to refine the peak set for training purposes below.

### ChromBPNet model training

To train the deep learning model, we utilized the suite of tools from ChromBPNet v2.5.0^46^ pipeline with default parameters on each consolidated BAM file from each cell type in the scATAC-seq dataset. The final input for model training consists of 183,485 parietal endoderm peaks identified above and a corresponding background peak set created using chrombpnet *prep nonpeaks*. We defined the chromosomal splits for training, testing, and validation, defined as “fold 0”, using chrombpnet *prep splits* on valid mm10 chromosomes (options:-tcr chr1 chr4 chr6-vcr chr10 chr17).

### ChromBPNet model predictions

To assess the functional relevance of each oligo sequence synthesized, we embedded each sequence at the center of 1,000 mouse genomic background sequences, which were deemed inaccessible for the specific cell type of interest, using a custom Python script. We then averaged the predictions across these background sequences to compute the marginal footprint for each sequence. The final prediction scores were normalized by the null value, which represents the activity of the background sequence alone.

### Design and cloning of CRE ortholog libraries for MPRA

Primers and plasmids are listed in **Table S2**.

To identify orthologs, we selected the maximal subsequences of five active CREs specifically expressed in parietal endoderm cells. These sequences were tiled into 270 bp windows with 5 bp steps, based on our subtiling data^38^, and symmetrically extended by 15 bp to obtain 300 bp sequences. Orthologous sequences were retrieved from 241 extant and reconstructed mammalian genomes using the *Cactus* alignment (2020v2) from the *Zoonomia* consortium^17^, facilitated by the Hal suite of tools^2^. Using HalLiftover (cactus-bin-v2.7.1), we mapped the mouse CRE subsequences (mm10) as queries against the genome intervals of all 241 species (options: –bedType 4 –noDupes 241-mammalian-2020v2.hal). The resulting possibly discontiguous output orthologous intervals were then merged with stitchHalFrags_v2.R, a custom script, requiring that the final interval was within 0.5 to 1.5x in length compared to the original query interval size. Sequences not meeting this size threshold were discarded from downstream analysis. In cases for which the intervals spanned different contigs, the sequences were also discarded to avoid complications. Sequences that failed to meet these criteria or contained undetermined bases (N) were discarded.

The resulting valid sequences were retrieved using bedtools getfasta (v2.29.2) from the corresponding target genomes, with only the highest-quality sequences included. All sequences were then aligned to the mouse reference using pairwise global alignment (Biostrings 2.62.0, pairwiseAlignment, options: type=“global”, gapOpening =-2, gapExtension =-8) to determine the correct orientation, with the orientation yielding the highest alignment score retained.

To construct the CRE ortholog library, the sequences were synthesized as 500 bp oligos (Twist Biosciences) with dial out primers for PCR amplification and isolation. Oligos were double stranded using three rounds of PCR amplification with KAPA2G Robust HotStart ReadyMix (Roche). A total of 5 ng of DNA input was used. For dial out PCR1, DNA was mixed with 12.5 µl KAPA2G Robust master mix, 1.25 μl 10 μM (F_Retrieval_Primer_48), 1.25 μl 10 μM (R_Retrieval_Primer_48), 0.25 μl of SYBr green 100×, and 8.75 μl water. Homology ends for Gibson assembly were appended through PCR2, with ∼5 ng of the eluate from PCR1 taken as input, 12.5 µl KAPA2G Robust master mix, 1.25 μl 10 μM oJBL684, 1.25 μl 10 μM oJBL685, 0.25 μl of SYBr green 100×, and 8.75 μl water. Barcode (BC) insertions for each element were appended through PCR3, with ∼5 ng of the eluate from PCR2 taken as input, 12.5 µl 2× KAPA2G Robust master mix, 1.25 μl 10 μM oJBL684, 1.25 μl 10 μM oJBL056, 0.25 μl of SYBr green 100×, and 8.75 μl water. Primer oJBL056 contains fifteen random Ns to serve as a molecular barcode (BC) for CRE-BC association. Each PCR step was amplified with tracking by qPCR with 1 min at 95 °C, and cycles up to the qPCR inflection point with 15 s at 95 °C, 15 s at 65 °C and 1 min at 72 °C. All PCR steps were cleaned up with DNA Clean & Concentrator (Zymo Research), eluted in 12 μl of water and quantified with a spectrophotometer (Nanodrop). The final product pool was used as inserts for a pooled Gibson assembly as described below.

The plasmid backbone (p001) was linearized using AgeI-HF and SbfI-HF (NEB) and ligated with the pooled PCR-amplified CRE orthologs via Gibson assembly. The assembly mixture was transformed (electroporation) into competent bacteria (C3020K; NEB). After bottlenecking to ensure library complexity of ∼150,000 clones, cultures were incubated overnight at 37°C, and plasmid DNA was purified the following day using a midiprep kit (Zymo Research). The resulting plasmid library was then used for the final subassembly step to associate BC to a CRE.

### Design and cloning of synthetic reconstitution libraries for MPRA

For each ancestral ortholog of the mouse CRE, we aligned the orthologs to their respective mouse reference sequences using pairwise global alignment (sequence_align ^74^; needleman_wunsch, options: match_score=1, mismatch_score=-1, indel_score=-1, gap = “_”). We then identified all mismatches and insertion/deletion (indel) events between the orthologous sequences and the mouse reference. These differences were introduced individually into the ancestral CRE sequences. Each mutated sequence was then evaluated using our deep learning model, which predicted the impact of each mutation on chromatin accessibility (see <u>ChromBPNet</u> <u>model predictions</u> section above). The mutation that predicted the greatest increase in accessibility was selected, added, and this process repeated in an iterative fashion until the full mouse CRE was reconstituted. As a control, random reconstitution trajectories were generated by introducing the same set of mouse-derived mutations in random order until the full mouse sequence was recovered. For each CRE, ten independent randomized trajectories were generated. All designs were created through a custom Python script.

Following this, we synthesized all members of each trajectory (Twist Biosciences). A total of 5 ng of DNA input was used. For dial out PCR1, DNA was mixed with 12.5 µl KAPA2G Robust master mix, 1.25 μl 10 μM (F_Retrieval_Primer_76 and F_Retrieval_Primer_11), 1.25 μl 10 μM (R_Retrieval_Primer_76 and R_Retrieval_Primer_11), 0.25 μl of SYBr green 100×, and 8.75 μl water. Homology ends for Gibson assembly were appended through PCR2, with ∼5 ng of the eluate from PCR1 taken as input, 12.5 µl KAPA2G Robust master mix, 1.25 μl 10 μM oJBL681, 1.25 μl 10 μM oJBL682, 0.25 μl of SYBr green 100×, and 8.75 μl water. Barcode (BC) insertions for each element were appended through PCR3, with ∼5 ng of the eluate from PCR2 taken as input, 12.5 µl 2× KAPA2G Robust master mix, 1.25 μl 10 μM oJBL681, 1.25 μl 10 μM oJBL056, 0.25 μl of SYBr green 100×, and 8.75 μl water. Primer oJBL056 contains fifteen random Ns to serve as a molecular barcode (BC) for CRE-BC association. Each PCR step was amplified with tracking by qPCR with 1 min at 95 °C, and cycles up to the qPCR inflection point with 15 s at 95 °C, 15 s at 65 °C and 1 min at 72 °C. All PCR steps were cleaned up with DNA Clean & Concentrator (Zymo Research), eluted in 12 μl of water and quantified with a spectrophotometer (Nanodrop). Plasmid library preparation was performed as described in the CRE ortholog library section, yielding a final library complexity of ∼180,000 clones.

### Design and cloning of ablation/enhancement trajectories for MPRA

For each 300 bp mouse CRE subsequence, we employed *in silico* mutagenesis to identify mutations predicted to either enhance or ablate chromatin accessibility. Using our deep learning model (see <u>ChromBPNet model</u> <u>predictions</u> section above), we predicted the effects of all possible mutations on accessibility and selected the most optimal mutation to either increase or decrease accessibility. This iterative process was repeated for 50 steps to accumulate sequential mutations. All designs were created through a custom Python script.

Following this, we synthesized all members of each trajectory (Twist Biosciences). PCR amplification and plasmid library preparation were performed as above (see <u>Design and cloning of synthetic reconstitution</u> <u>libraries for MPRA</u> section), yielding a final library complexity of ∼50,000 clones. Dial out primer sequences used for amplification were F_Retrieval_Primer_65 and R_Retrieval_Primer_65.

### CRE-BC subassembly

To associate barcodes to CRE, PCR amplification was performed from the plasmid libraries to generate the CRE-BC libraries. The reaction consisted of 12.5 µl KAPA2G Robust master mix, 1.25 μl 10 μM oJBL060 (appending Nextera P7 adapter), 1.25 μl 10 μM custom-indexed P5 primers (NextP5_index1-9), 0.25 μl of SYBr green 100×, and 8.75 μl water. PCR step was amplified with tracking by qPCR with 1 min at 95 °C, and cycles up to the qPCR inflection point with 15 s at 95 °C, 15 s at 65 °C and 1 min at 72 °C. The resulting amplified libraries are purified by 1.0× Ampure XP beads (Beckman Coulter).

Sequencing libraries were pooled and paired-end sequenced on the NextSeq2000 with the following primers and cycle numbers: read1 (CRE forward): 159 cycles, primer oJBL681/oJBL684; index1 (BC read): 15 cycles, primer oJBL061; read2 (CRE reverse): 158 cycles, primer oJBL683/oJBL686; index2 (sample index): 6 cycles, primer oJBL064. For libraries with larger amplicons, cycle numbers were adjusted: read1: 401 cycles; index1: 15 cycles; read2: 212 cycles; index2: 10 cycles.

CRE reads were preprocessed to remove adapter reads and low quality base calling using *trim_galore*^75^ (default parameters for paired-end reads). After trimming, forward and reverse CRE reads were then joined and error-corrected with PEAR^76^ (small amplicons: options-v 4; larger amplicons: options-v 20). Next, CRE reads were mapped to their respective reference libraries using *bowtie2* ^77^ (v2.5.3) and BC reads were merged and tallied for each mapped CRE using custom Python and R scripts. Following this, BCs were filtered based on read count to remove low-abundance barcodes from the CRE-BC association. The resulting CRE-BC association dictionaries were utilized for downstream analyses, which included processing valid barcode counts for bulk MPRA.

### MPRA library transfection and collection

Three biological replicates of 2-6 M PYS-2 cells were transfected episomally using Lipofectamine 2000 (Thermo Fisher, cat. no. 11668030, Gibco Opti-MEM cat. no. 31985) with 4 µg of reporter plasmid per replicate. Positive (plasmid 002) and negative (plasmid 003) control plasmids were transfected at matched concentrations to monitor transfection efficiency and to assess the duration of episomal plasmid expression following transfection. Cells were washed with phosphate-buffered saline (PBS) with medium changes the next day, and cells passaged as usual thereafter. After 2 days, PYS-2 cells were lifted off plates with 0.05% trypsin, washed once with PBS, and resuspended in 80% ice-cold methanol, and placed at −80 °C until further processing.

### Bulk MPRA library preparation

Genomic DNA and RNA was extracted from methanol fixed cells using the All-prep DNA/RNA mini kit (Qiagen) following the manufacturer’s instructions. MPRA amplicon libraries from DNA were generated in two steps of PCR amplification with KAPA2G Robust HotStart ReadyMix (Roche). A total of 0.5–1 µg of genomic DNA input was used. For low-cycle number PCR1, gDNA was mixed with 12.5 µl KAPA2G Robust master mix, 1.25 μl 10 μM oJBL790, and 1.25 μl 10 μM oJBL789. Cycling parameters: 1 min at 95 °C, and four cycles of 15 s at 95 °C, 15 s at 65 °C and 30 s at 72 °C, followed by 4 °C hold. Primer oJBL789 contains ten random Ns to serve as a pseudo-UMI (hereafter referred to as UMIs for brevity) to correct for PCR jackpotting. Reactions were cleaned up with Ampure XP beads at 1×, and eluted in 12 uL of water. Illumina adapters and sequencing indices were appended through PCR2, with 5 μl of the eluate from PCR1 taken as input, and 7.5 μl KAPA2G Robust master mix, 0.75 μl 100× SYBr green, 1.25 μl 10 μM oJBL076, and 1.25 μl 10 μM Nextera P7 indexed primers. Libraries were amplified with tracking by qPCR with 1 min at 95 °C, and cycles up to the qPCR inflection point with 15 s at 95 °C, 15 s at 65 °C and 1 min at 72 °C. Libraries were then cleaned up with Ampure XP beads at 1×.

Amplicon libraries for RNA were obtained by first DNase-treating RNA (5 μg RNA, 2.25 μl TURBO DNase (Thermo Fisher), and 1.875 μl 10× buffer, incubated at 37 °C for 60 min, followed by 3.8 μl of TURBO DNase inactivation buffer. One microgram of DNase-treated RNA was then taken to reverse transcription. Briefly, 11 μl (500 ng μl−1) RNA was mixed with 2 μl 2 μM oJBL789, incubated at 65 °C for 5 min and placed on ice. Seventeen microliters of reverse transcription master mix was then added (4 μl 5× FS buffer, 1 μl 0.1 M dithiothreitol, 1 μl 10 mM dNTP mix, and 1 μl SSIII (Thermo Fisher)), and the reaction was incubated at 55 °C for 60 min, followed by 80 °C for 10 min. Half of the reverse transcription reaction was then directly amplified for PCR1 (12.5 μl KAPA2G Robust master mix, 1.25 μl 10 μM oJBL790, and 1.25 μl 10 μM Nextera P7 indexed primers), with cycling parameters of 1 min at 95 °C, and four cycles of 15 s at 95 °C, 15 s at 65 °C and 30 s at 72 °C, followed by 4 °C hold. Reactions were cleaned up with Ampure XP beads at 1.5 ×, and eluted in 12 μl of water. PCR2 proceeded as for libraries prepared from genomic DNA, with oJBL077 and oJBL076, and reactions were stopped at inflexion point from qPCR tracking. Libraries were then cleaned up with Ampure XP beads at 1×.

Final libraries were quantified with TapeStation D1000 HS (Agilent) for final quality assessment, and adjusted to final 2 nM on the basis of the TapeStation quantification. Libraries were pooled and paired-end sequenced on NextSeq2000 with the following primers and cycle numbers: read1 (BC forward): 15 cycles, primer oJBL072; index1 (sample index): 10 cycles, primer oJBL760; read2 (UMI): 10 cycles, primer oJBL761; index2 (BC reverse): 15 cycles, primer oJBL074.

### Bulk MPRA data processing and quantification

Sequencing data were demultiplexed using bcl2fastq. Forward and reverse mBC reads were joined and error corrected with PEAR v0.9.11^76^ (options:-v 15-m 15-n 15-t 15). Using custom Python and R scripts, successfully assembled barcode reads were combined with UMI reads, BC–UMI pairs were counted, and the read and UMI counts per BC were determined. The read and UMI counts for the BC present in the reporter pools (determined a priori; see <u>CRE-BC subassembly</u> section above) were collected as comparison to the bulk measurements.

Expression for each BC from the UMI counts table was computed as follows. First, the total UMI per sample (per cell line, replicate, and batch) to the BC in our list was determined for both RNA- and DNA-derived libraries. Each BC UMI count was then normalized by the summed of counts in its respective sample type (DNA and RNA). The normalized RNA UMI count was then divided by the normalized DNA UMI count (1% winsorized summed RNA over DNA UMI count), to generate the bulk MPRA-derived estimate of expression per BC.

Statistical analyses for individual elements were performed using *BCalm*^78^ R package, comparing activity estimates to those of standard negative control sequences.

### TF binding affinity calling

To assess transcription factor binding affinity for each sequence in the dataset, we utilized ProBound v1.0^79^ with default settings:(loadMotifCentralModel(TFID).addNScoring().inputTXT(seq.txt).bindingModeScores(/dev/stdout,pr ofile) for the following TFBSs: Gata4, Gata6, Foxa2, Sox17, and Jun_Atf3 (TFID: 16735, 17095, 17084, 10629, 15946). For each sequence, a sliding window corresponding to the *k*-mer size of the TFBS was applied at every base pair, allowing us to measure the binding affinity at each subsequence, considering both the forward and reverse orientations.

Subsequent to the binding affinity calculations, we normalized the TF binding affinities by comparing them to the maximum affinity observed in a randomly generated set of 1 million *k*-mers (affinity > q99.99). This normalization ensures that the TFBS binding affinities are consistent across sequences. To infer the orientation of each TFBS, we compared the normalized binding affinities in both orientations. If the forward orientation affinity was greater than or equal to the reverse orientation, the TFBS was classified as forward. Otherwise, it was classified as reverse. The final binding affinity for each TFBS was assigned based on the higher of the two normalized affinities (forward or reverse orientation). Given the redundancy between Gata4 and Gata6 TFBSs, we grouped the binding affinities of these two TFs under the label Gata4/6, using the maximal normalized binding affinity observed for each. Finally, only sequences with a normalized binding affinity greater than 0.05 were considered as putative TFBSs.

### Functional TFBS alignment

For each set of CREs, we performed multiple sequence alignment (MSA) using MAFFT v7.505^80^ (options: –adjustdirection). The alignment was then mapped to the corresponding mouse reference sequence, enabling the linking of MSA positions to the reference positions. This mapping provides a direct connection between the aligned sequences and the reference mouse sequence, facilitating the identification of functional mouse TFBS within each orthologous CRE identified in our saturation mutagenesis data^38^.

Following the TF binding affinity analysis (see <u>TF binding affinity calling</u> above), we identified putative TFBSs by overlapping the sites with the functional mouse TFBSs using the GenomicRanges *findOverlap* function. All subsequent visualizations of TFBS alignments were generated using the putative TFBSs, mapped to each sequence along the MSA positions.

### Evolutionary lineage tracing

To trace the lineage from the first mammalian ancestor to its respective extant species, we used the Newick tree format provided by *Cactus*^2^, which includes the 241 species and their reconstructed ancestral genomes. Phylogenetic trees were generated and visualized using the *ggtree* package^81^. Evolutionary lineage tracing was performed using the ape package, specifically the *nodepath* function, to identify all ancestral nodes leading to each extant species for further analysis.

## SUPPLEMENTARY TABLES

**Table S1**: Extant and ancestral ortholog recovery per CRE.

**Table S2**: Primers and plasmids used in experiments.

## SUPPLEMENTARY FIGURES 1-11

**Supplementary Figure 1.**
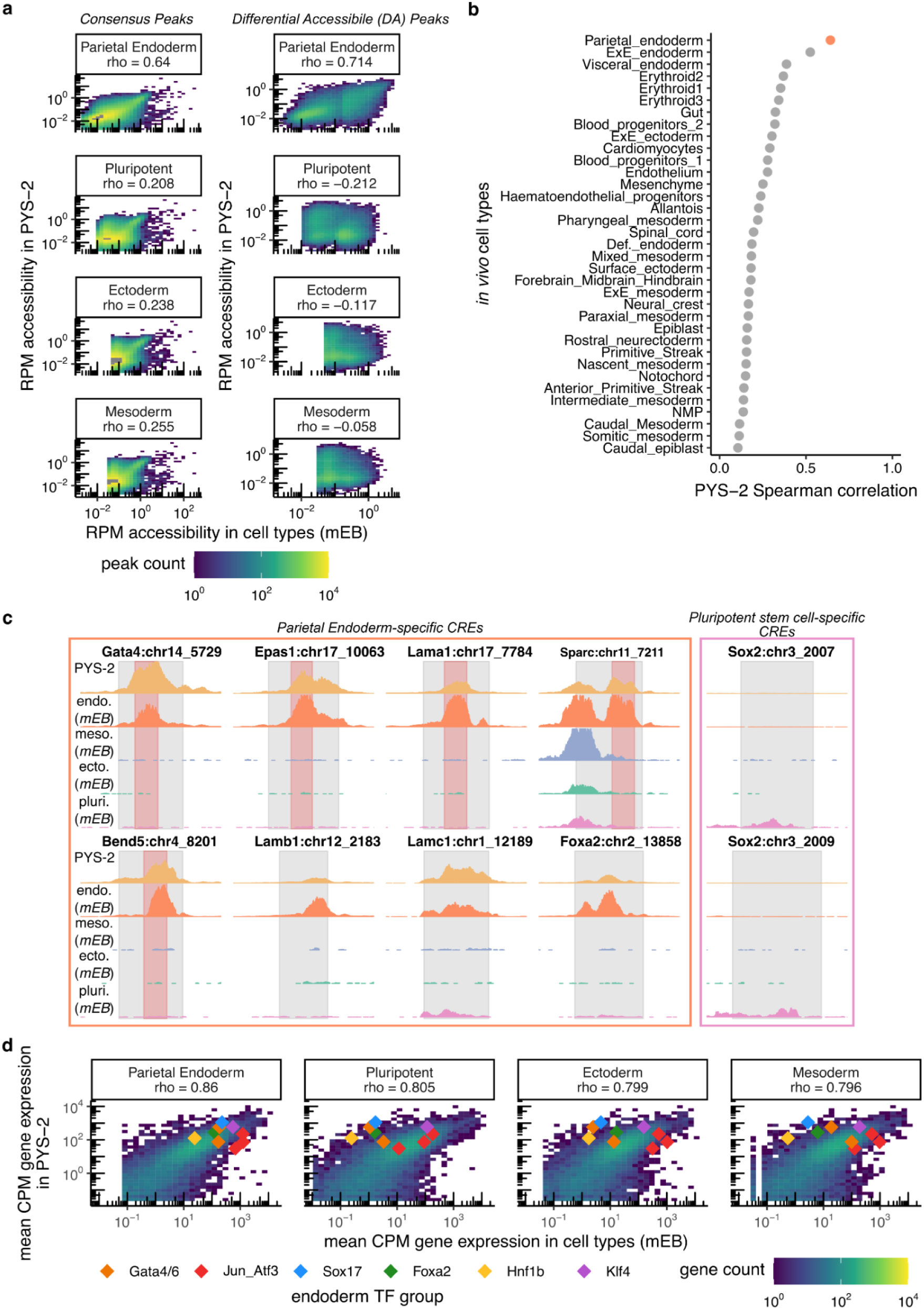
Evaluation of the PYS-2 cell line as a surrogate for the parietal endoderm lineage. **a)** Comparison of normalized read counts (RPM) at consensus peaks (left) or differentially accessible peaks (right) in ATAC-seq data from mEB cell types^34^ (pseudobulk; x-axes) vs. PYS-2 (bulk; y-axes). Consensus peaks are those called in both PYS-2 and mEB cells (*n* = 426,803). Differential peaks are those called as differential across mEB cell types (*n* = 56,795). Correlations (Spearman’s *ρ*) are based only on peaks with RPM > 0 in both cell types being compared. **b)** Correlations (Spearman’s *ρ*) for comparison of RPM at consensus peaks (*n* = 206,973) in ATAC-seq data from *in vivo* mouse cell types^37^ (pseudobulk) vs. PYS-2 (bulk). **c)** Chromatin accessibility tracks for eight parietal endoderm-specific CREs (left) and two pluripotent stem cell-specific CREs (right) validated by scQer^34^, in either PYS2 cells (top track) or mEB germ layers (bottom four tracks). Shaded gray regions indicate the genomic coordinates of the original full-length CREs as validated by scQer^34^ (520 bp to 1.7 kb in length). Shaded red regions correspond to coordinates of 300-bp maximum activity tile used for downstream experiments for 5 of the CREs. **d)** Comparison of mean gene expression counts (CPM) in RNA-seq data from mEB cell types (pseudobulk; x-axes) vs. PYS-2 (bulk; y-axes). Correlations (Spearman’s *ρ*) are based on the log-transformed CPM expression profiles.

**Supplementary Figure 2.**
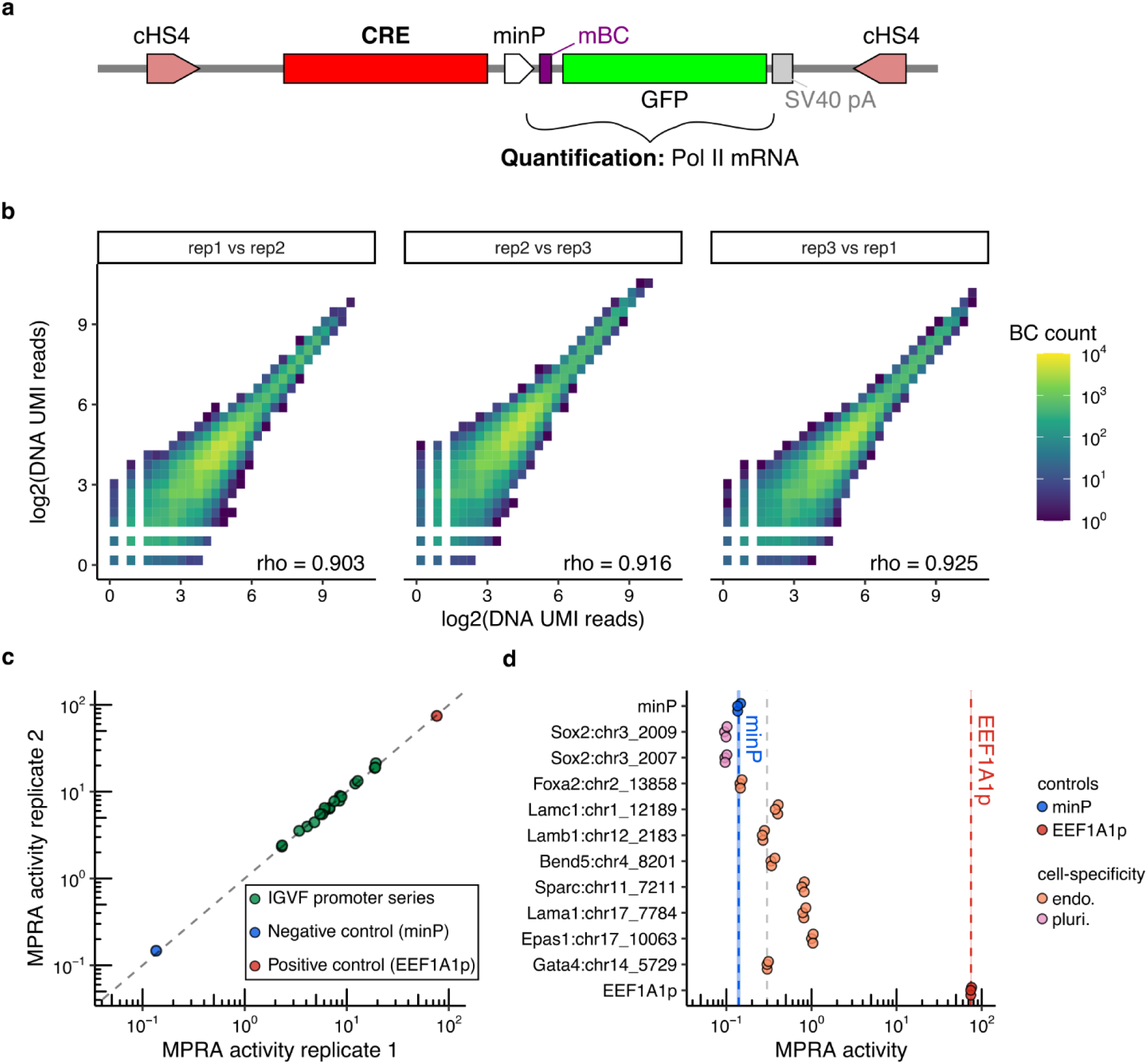
**Quality control of MPRA-based activity profiling in PYS-2 cells**. **a)** Schematic of the MPRA reporter cassette used for transient transfection. A library of CRE is cloned to a position upstream of a minimal promoter (minP) driving a reporter gene containing a barcoded 5′ untranslated region (mBC) and a GFP open reading frame, followed by the SV40 polyadenylation signal (SV40 pA). These elements are all flanked by convergent insulators (core chicken hypersensitive site-4 from beta-globin locus, cHS4^82^). **b)** Scatterplot comparing biological replicates with respect to their log2-transformed UMI read counts for DNA-derived barcodes, each associated with a specific CRE being tested. Replicates here are independent transient transfections of the same MPRA library into PYS-2 cells (≥5 million cells transiently transfected per replicate; cells fixed 48 hours post-transfection). Points represent UMI read counts per barcode for the original parietal endoderm CREs (avg. replicate barcode recovery per element = 8781) and promoter series (avg. replicate barcode recovery per element = 105, 8799 and 4242 for IGVF, minP and EEF1A1p, respectively). Correlations (Spearman’s *ρ*) were calculated by comparing the log_2_-transformed UMI read counts of pairs of replicates. **c)** Mean MPRA barcode activity (UMI read counts from RNA/UMI read counts from DNA) of promoters in transfection replicate 1 vs. 2. Promoters include the internal IGVF promoter series (green), minimal (minP, blue), and EEF1A1 promoter (red). **d)** MPRA activities (x-axis) of ten full-length scQer-nominated cell type-specific CREs^34^, minimal promoter (minP) and EEF1A1 promoter (rows), all in PYS-2 cells. Each point represents the measured activity of a CRE in a transfection replicate. Blue dotted line and shading indicates the MPRA activity mean and standard deviation of the background control (minP), while the grey dotted line represents the 2-fold activity threshold above the background control.

**Supplementary Figure 3.**
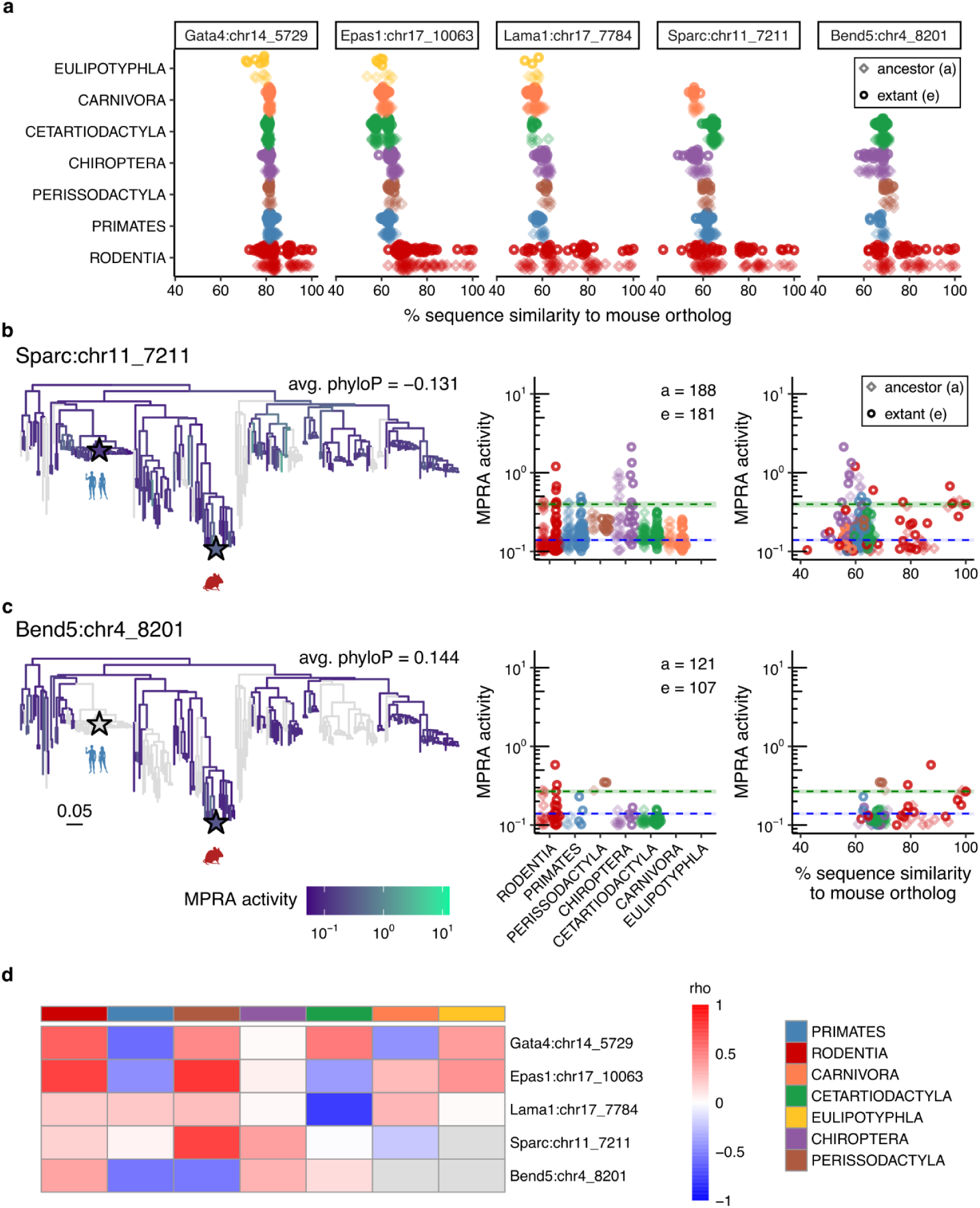
Sequence analysis of mammalian CRE orthologs. **a)** Sequence identity (%) of the CRE orthologs with the endogenous mouse CRE across taxonomic orders. Lighter colors represent ancestral orthologs, and darker colors represent extant sequences. **b-c)** Left: Mammalian phylogenetic tree onto which we have projected the MPRA-measured functional activity of ancestral or extant orthologs of the *Sparc* **(b)** and *Bend5* **(c)** proximal CREs in PYS-2 cells. MPRA activities represent normalized RNA/DNA ratios. Grey branches denote species for which the ortholog was either not present in the *Cactus* alignments/reconstructions or not recovered in the assay. Colored stars denote the positions/activities of humans and mice. The mean phyloP conservation score for bases in the mouse ortholog is also shown. A scale bar at the bottom indicates branch length in units of substitution per site. Middle: MPRA activity (*y*-axes) broken out by taxonomic order (*x-*axes, colors). Right: MPRA activity (*y*-axes) as a function of sequence divergence from mouse (*x-*axes, colors). In both middle and right plots, the green and blue dotted lines correspond to activity of the mouse ortholog and minP background control, respectively. Lighter hues are used for ancestral species, and darker hues for extant species. **d)** Heatmap of correlations (Spearman’s *ρ*) between MPRA measurements vs. sequence similarity to the extant mouse ortholog for each of the 5 CREs (rows), broken out by taxonomic order (columns). Grey indicates no available measurement.

**Supplementary Figure 4.**
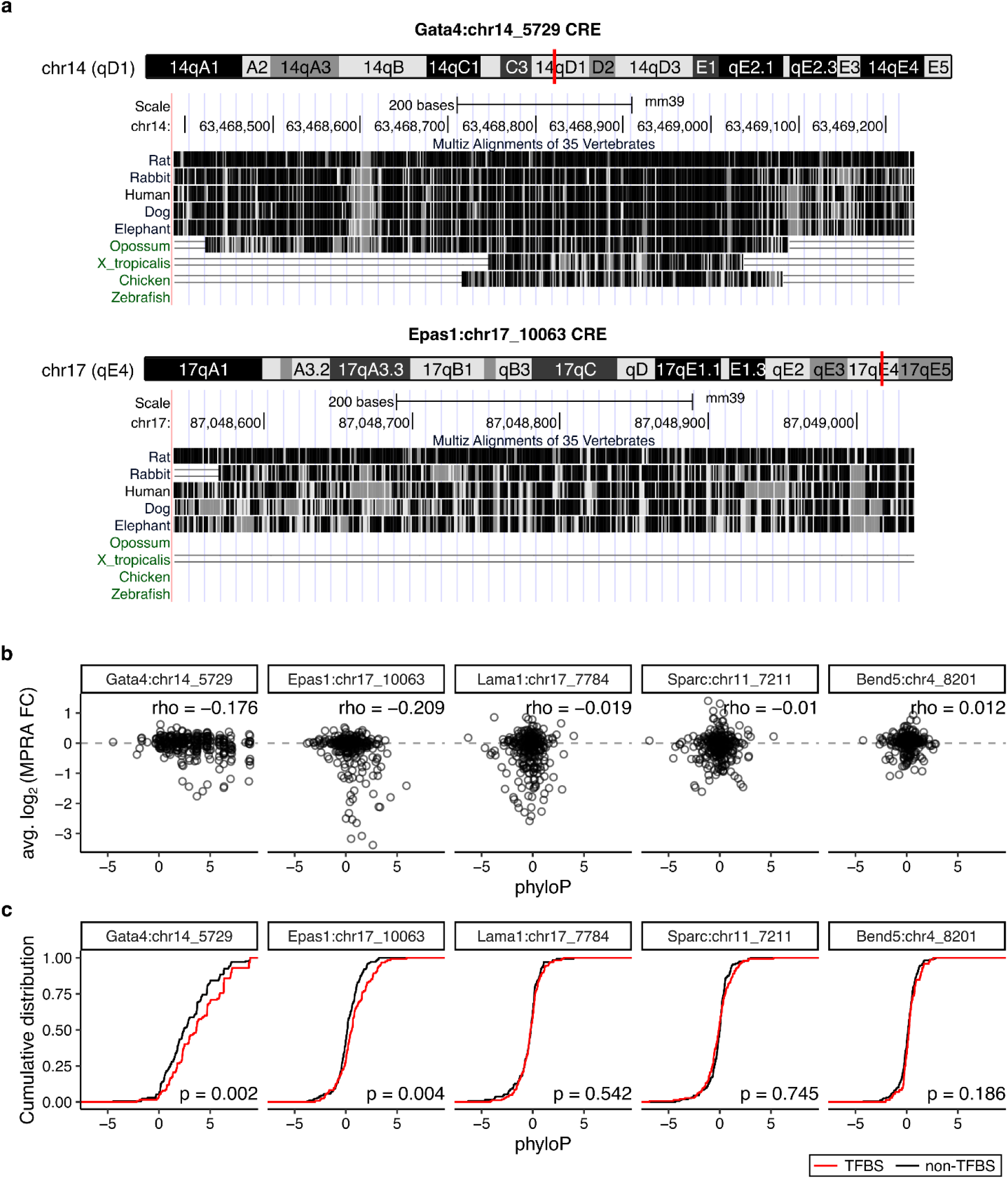
Conservation vs. saturation mutagenesis for parietal endoderm mouse CREs. **a)** UCSC Multiz alignment of 60 vertebrates showing the regions containing the *Gata4* (top; chr14:63,468,388-63,469,232) and *Epas1* (bottom; chr17:87,048,296-87,049,290) CREs in the mouse genome (GRCm39/mm39). Conservation levels are indicated by shading, with darker regions corresponding to higher conservation as scored by phastCons. Placental mammals are highlighted in blue, and non-placental vertebrates in green. **b)** Scatter plot showing the lack of correlation between phyloP scores and the functional consequences of single nucleotide substitution across the CRE as measured in PYS-2 cells^38^. Correlations (Spearman’s *ρ*) were calculated between phyloP scores and mean log_2_ fold-changes in MPRA activity. **c)** Cumulative distributions of phyloP scores (calculated from Cactus 241 mammalian species) for individual nucleotides within the CRE subsequences, comparing positions overlapping functional mouse TFBSs (red) to non-TFBS positions (black). P-values were computed to test for differences in phyloP distributions between TFBS sequences and non-TFBS sequences for each CRE using the Wilcoxon rank sum test.

**Supplementary Figure 5.**
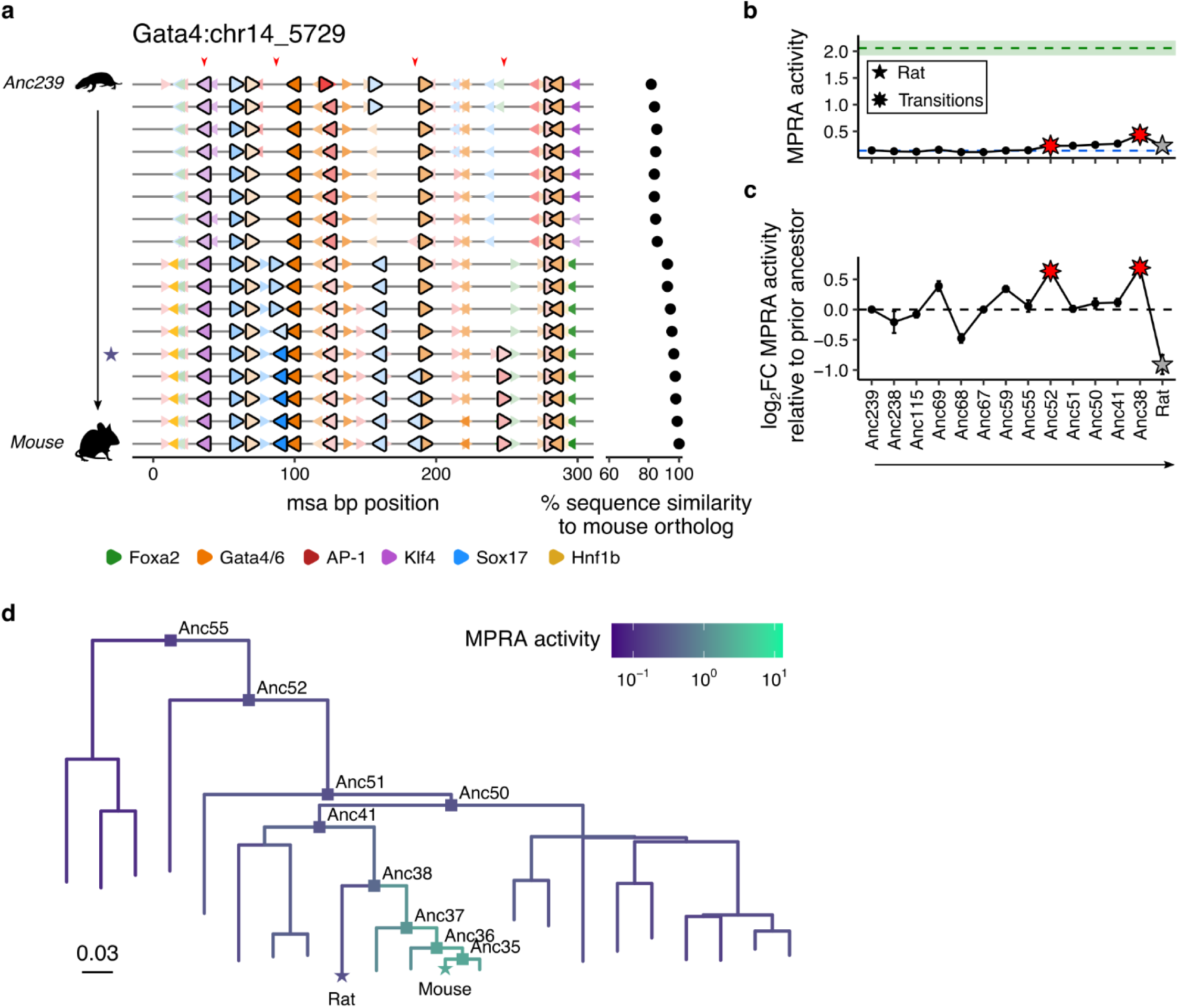
Functional evolution of the *Gata4* CRE in Rodentia. **a)** Positions of predicted TFBS for each ortholog of the *Gata4* CRE along the evolutionary trajectory leading to *M. musculus*, with the subset that are functional by saturation mutagenesis MPRA assigned larger triangles with black outlines. Hue color for these predicted TFBS corresponds to predicted affinity strength, normalized across the set of sequences shown. The purple star at the left indicates the common ancestor of mouse (*M. musculus*) and rat (*R. norvegicus*). The red arrowheads at the top indicate the four TFBS that are highlighted in **Fig. 2f-g** and discussed in the main text. At the right, % sequence similarity of each ancestral ortholog to the extant mouse ortholog is shown. **b-c)** Change in MPRA activity profile (top) and the log_2_ fold-changes in MPRA activity relative to the immediately prior ancestor (bottom) along the evolutionary trajectory of the *Gata4* CRE leading to *R. norvegicus*. Error bars indicate the standard error across three experimental replicates. **d)** Same as **Fig. 1c**, but zooming in on a Rodentia subclade rooted at Anc55 and colored by the measured MPRA activity of *Gata4* CRE orthologs. Squares highlight ancestral nodes along the mouse and rat lineage. Stars indicate extant mouse and rat. A scale bar at the bottom indicates branch length in units of substitution per site.

**Supplementary Figure 6.**
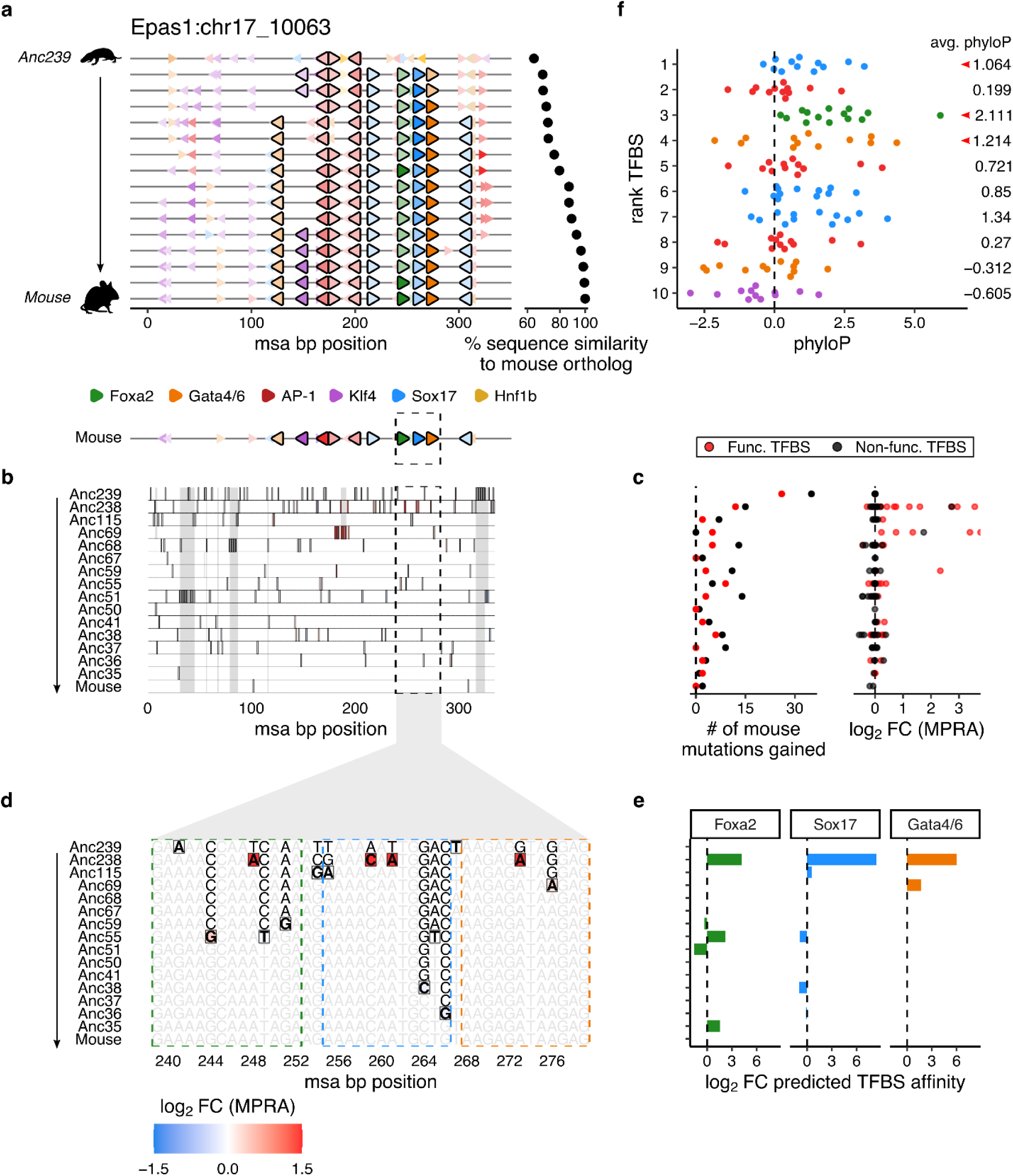
Evolution of a conserved *cis*-regulatory module in the *Epas1* CRE. **a)** Predicted TFBS positions for each ortholog of the *Epas1* CRE along the evolutionary path from Anc239 to *M. musculus*. TFBSs that are functional by saturation-mutagenesis MPRA are indicated by larger triangles with black outlines. Hues correspond to predicted TFBS affinities, normalized across the set of sequences shown. At right, the percent sequence similarity to the extant mouse ortholog is plotted. **b)** MSA of all *Epas1* CRE sequences along the evolutionary path from Anc239 to *M. musculus*. At top, predicted TFBS positions for the extant mouse ortholog are shown. Black-bordered tiles correspond to derived mutations relative to the immediately prior ancestor and are colored by log₂ fold-changes in saturation-mutagenesis MPRA activity^38^ (color scale at bottom left of figure; measured in mouse background). Gray columns mark derived indels not assayed by MPRA. Black dashed box denotes the conserved triplet module, mutations within which are further detailed in panels **d**-**e**. **c)** Number (left) and distribution of log₂ fold-changes in MPRA activity (right) for derived mouse mutations appearing at each step along the evolutionary path to the *M. musculus Epas1* CRE, partitioned by whether mutations lie within (red) or outside (black) a functional mouse TFBS. **d)** Nucleotide-resolution view of subregion of MSA shown in panel **b**, highlighting mutations within the conserved triplet module. Black-bordered tiles correspond to derived mutations relative to the immediately prior ancestor and are colored by log₂ fold-changes in saturation-mutagenesis MPRA activity^38^ (color scale at bottom left of figure; measured in mouse background). Gray columns mark derived indels not assayed by MPRA. Colored dashed boxes show the positions of the three TFBS—Foxa2 (green), Sox17 (cyan), and Gata4/6 (orange)—that comprise the conserved module. **e)** Log₂ fold-change in predicted TFBS affinity across evolutionary steps for the three conserved TFBSs highlighted in panel **d**. Key mutations from Anc239 → Anc238 include: T→A (pos 248), associated with an 18-fold gain in predicted Foxa2 affinity and 6.5-fold gain in measured MPRA activity; A→C (pos 259) and T→A (pos 261), associated with a 338-fold gain in predicted Sox17 affinity (together) and 7.5- and 12-fold gains in measured MPRA activity (respectively); and G→A (pos 273), associated with a 65-fold increase in predicted Gata4/6 affinity and 3.0-fold gain in measured MPRA activity. **f)** PhyloP scores for the 10 functional mouse TFBSs identified by a saturation mutagenesis MPRA of the *Epas1*:chr17_10063 CRE. Each row represents a functional TFBS ranked by the magnitude of its disruption on enhancer activity (top = highest). Each point denotes the phyloP score of one base-pair within the TFBS represented on that row. At right, the mean phyloP score for each TFBS is listed; red arrows denote the three TFBSs comprising the conserved heterotypic triplet module.

**Supplementary Figure 7.**
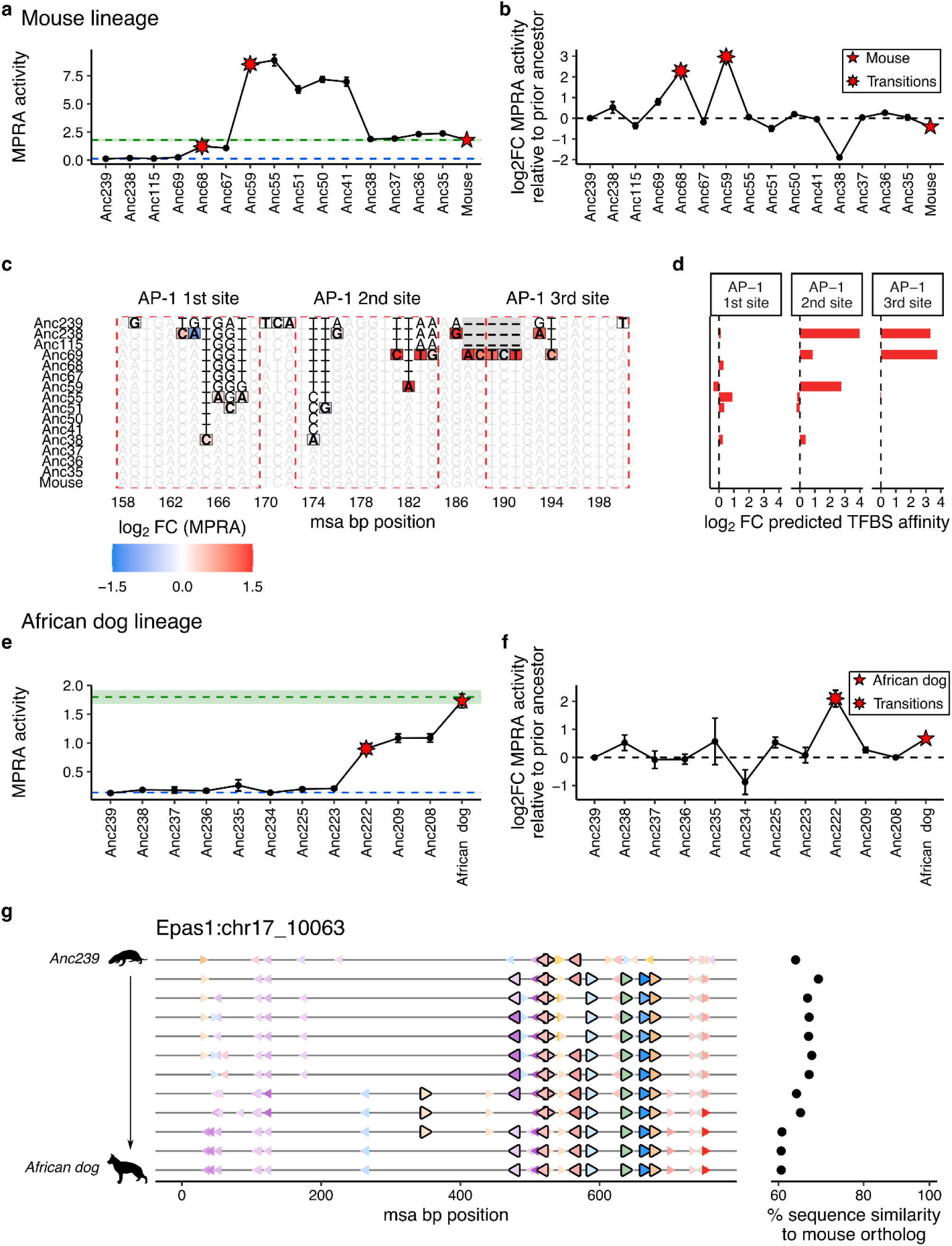
Evolution of species-specific AP-1 sites in mouse and African dog *Epas1* CRE ortholog. a–b) MPRA activity profiles (left) and log₂ fold-changes in MPRA activity relative to the immediately prior ancestor (right) of the *Epas1* CRE along the evolutionary path from Anc239 to *M. musculus*. Error bars indicate the standard error across three biological replicates. MSA of all *Epas1* CRE sequences along the evolutionary path from Anc239 to *M. musculus*. **c)** Nucleotide-resolution view of a subregion of MSA shown in **Fig. 4a** corresponding to a portion of the homotypic AP-1 module. Black-bordered tiles correspond to derived mutations relative to the immediately prior ancestor and are colored by log₂ fold-changes in saturation-mutagenesis MPRA activity^38^ (color scale at bottom left; measured in mouse background). Gray columns mark derived indels not assayed by MPRA. Dashed red boxes show the positions of the three AP-1 TFBS that comprise the homotypic cluster. **d)** Log_2_ fold-change in predicted TFBS affinity across evolutionary steps for the three AP-1 TFBSs highlighted in panel **c**, with matching rows as well. **e–f)** MPRA activity profiles (left) and log₂ fold-changes in MPRA activity relative to the immediately prior ancestor (right) of the *Epas1* CRE along the evolutionary path from Anc239 to *L. pictus* (African wild dog). Error bars indicate the standard error across three biological replicates. **g)** Predicted TFBS positions for each ortholog of the *Epas1* CRE along the evolutionary path from Anc239 to *L. pictus* (African wild dog). TFBSs that are functional by saturation-mutagenesis MPRA are indicated by larger triangles with black outlines. Hues correspond to predicted TFBS affinities, normalized across the set of sequences shown. At right, the percent sequence similarity to the extant mouse ortholog is plotted.

**Supplementary Figure 8.**
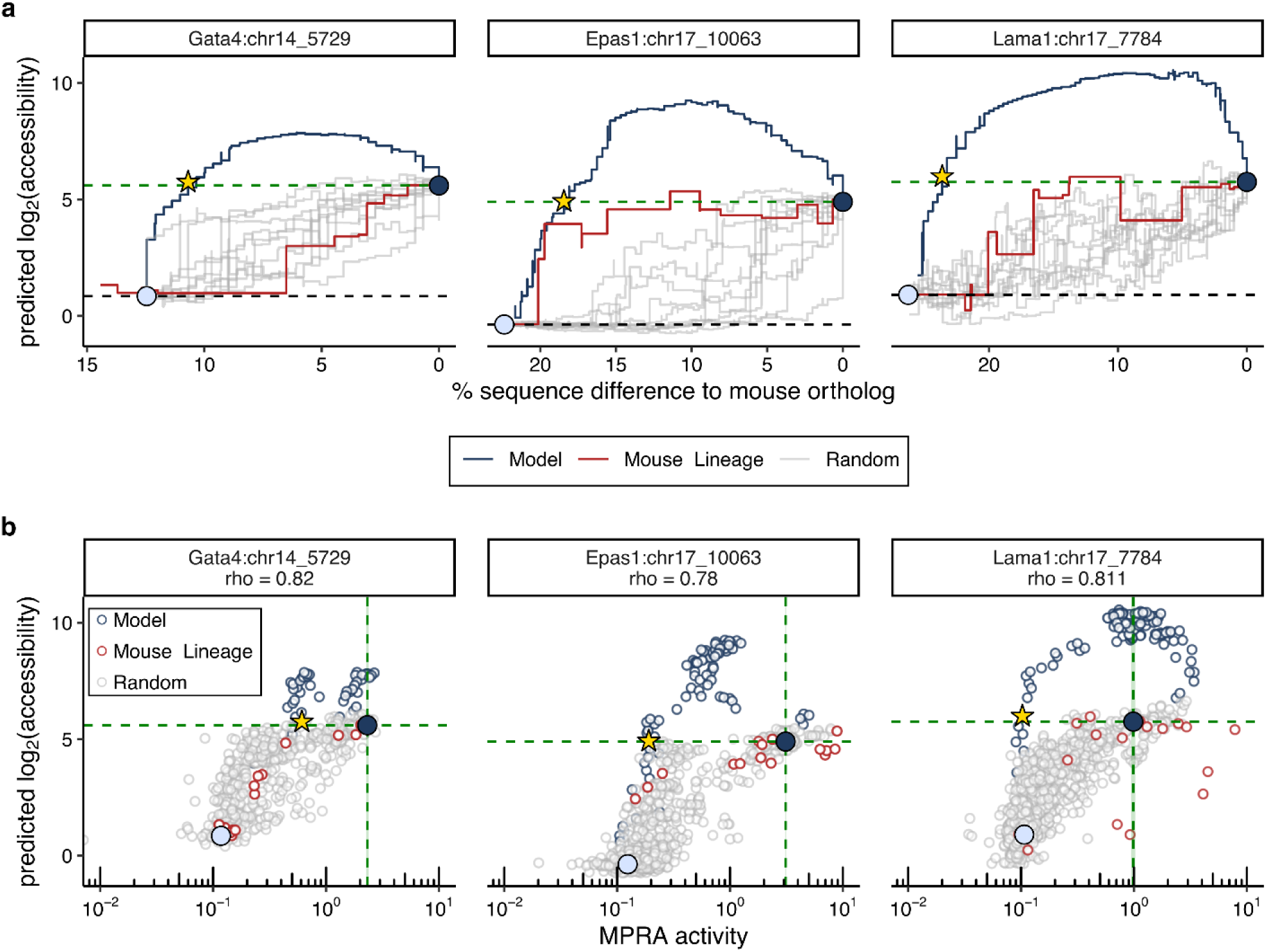
Comparison of model-predicted chromatin accessibility and experimentally measured enhancer activity during synthetic reconstitution. **a)** Model predicted log_2_ accessibility (y-axis) versus sequence divergence (x-axis) for the reconstitution trajectories and the mouse evolutionary lineage of the *Gata4*, *Epas1*, and *Lama1* CREs (from top to bottom). Model-optimized, random-order, and phylogeny-inferred evolutionary trajectories are shown in navy, gray, and red, respectively. Green and gray horizontal dotted lines indicate the predicted accessibility of the extant mouse ortholog and the inferred common mammalian ancestral ortholog, respectively. **b)** Scatterplots comparing MPRA activity (x-axis) and ChromBPNet-predicted log2 accessibility (y-axis) for all intermediate sequences across reconstitution trajectories. Points are colored by trajectory type (model-optimized, navy; random, gray; evolutionary lineage, red). Spearman correlation coefficients are shown for each CRE at the top. Large circles denote the initial (light blue) and final (navy) intermediates for all trajectories, and stars denote the earliest intermediate reaching or exceeding the predicted accessibility of the endogenous mouse CRE.

**Supplementary Figure 9.**
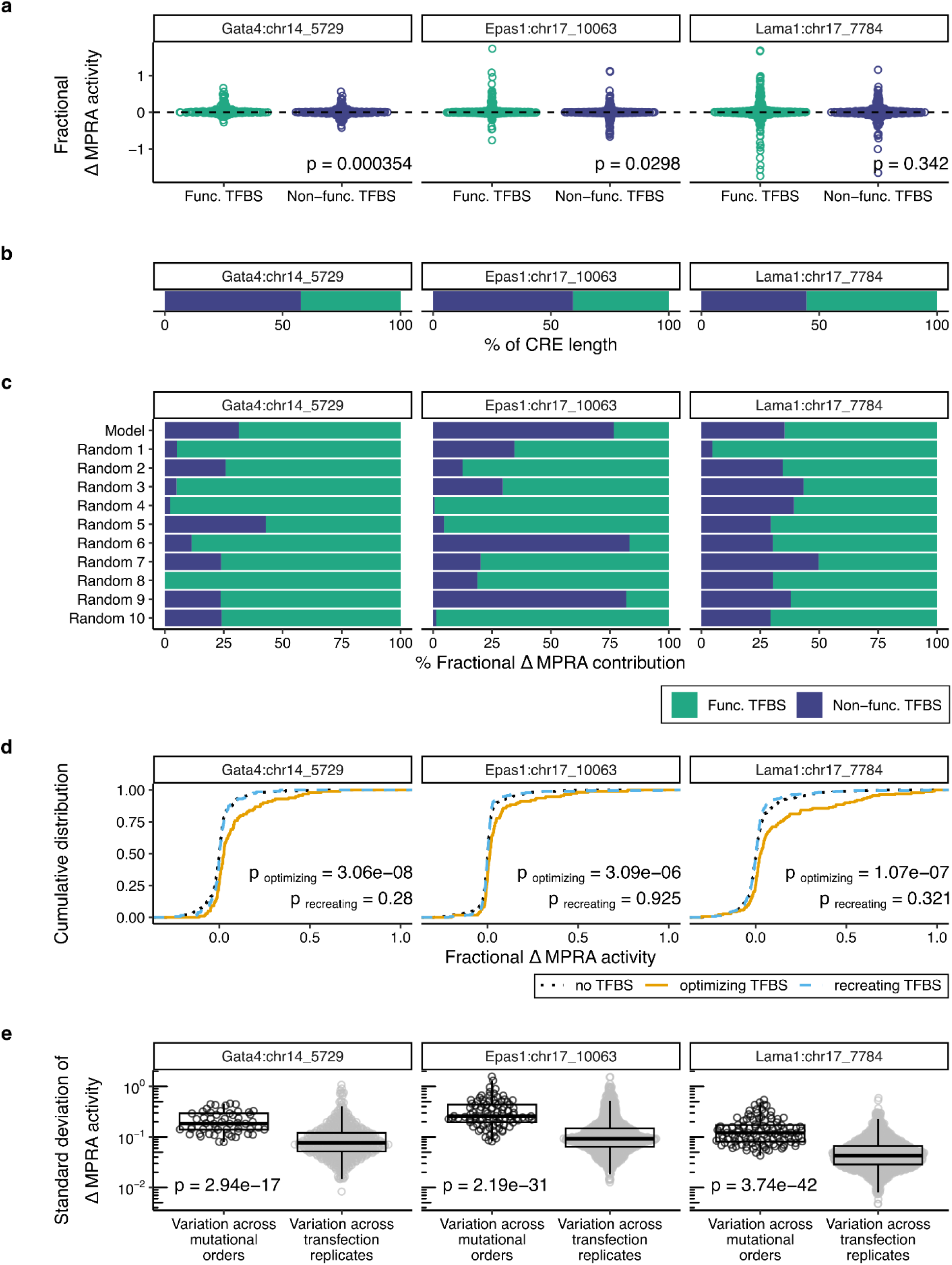
Contribution of TFBS-associated mutations and epistasis to enhancer reconstitution trajectories. **a)** Fractional changes in MPRA activity associated with individual mutations across all reconstitution trajectories, stratified into mutations overlapping functional TFBS (green) or non-TFBS (purple) positions, in the extant mouse CRE. P-values report Wilcoxon rank-sum tests comparing TFBS-overlapping and non-TFBS mutations for each CRE. **b)** Overall fraction of each CRE sequence overlapping functional TFBSs versus non-TFBS positions. **c)** Proportion of total fractional MPRA activity recovery attributable to mutations overlapping functional TFBSs (green) or non-TFBS (purple) positions, across all reconstitution trajectories for each CRE. **d)** Cumulative distributions of fractional MPRA activity changes grouped by mutation class: non-TFBS positions (black, dotted), TFBS-recreating mutations (cyan, dashed), and TFBS affinity-optimizing mutations (gold, solid). P-values indicate Wilcoxon rank-sum tests comparing TFBS-recreating or TFBS-optimizing mutations against mutations at non-TFBS positions, computed separately for each CRE. **e)** Comparison of order-dependent variability in mutational effects to experimental noise. Shown is the standard deviation of MPRA activity changes for each mutation across reconstitution trajectories (black; n = 11 trajectories) and across biological replicates within a trajectory (gray; n = 6 biological replicates per trajectory). Each point represents a single mutation. Box-and-whisker plots indicate the median, interquartile range, and whiskers extending to 1.5× IQR. P-values report Wilcoxon rank-sum tests comparing trajectory-level and replicate-level variability for each CRE.

**Supplementary Figure 10.**
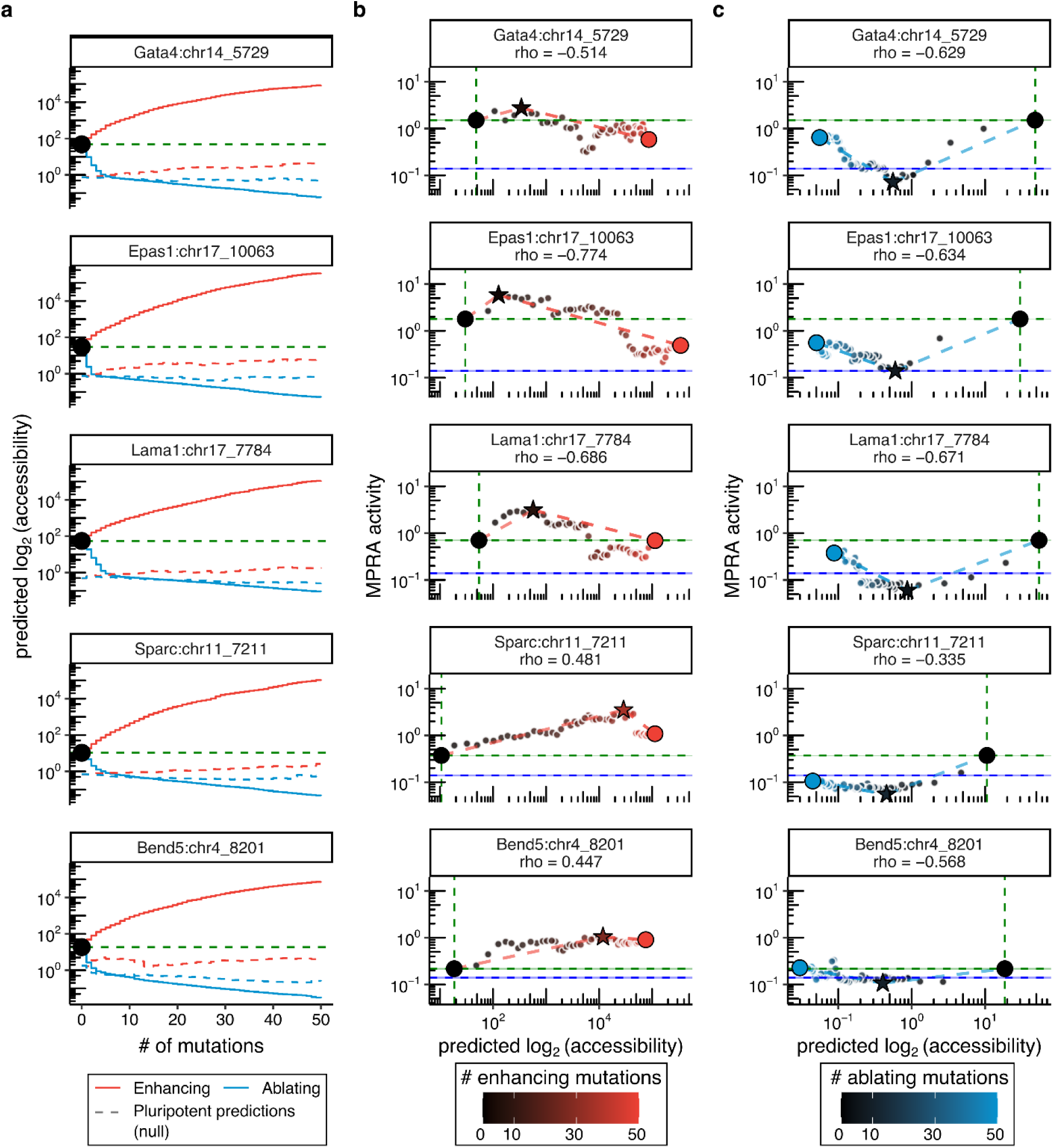
Comparison of predictions vs. observations upon model-guided enhancement or ablation of activity of endogenous parietal endoderm CREs. **a)** ChromBPNet-predicted log₂ chromatin accessibility (y-axis) across sequential gradient steps (x-axis) for model-guided modulation trajectories of the *Gata4*, *Epas1*, *Lama1*, *Sparc*, and *Bend5* CREs (top to bottom). Trajectories optimized for enhancement (red) or ablation (blue) are shown. Black circles denote the predicted accessibility of the endogenous mouse sequence, and green dotted lines indicate the same reference level across steps. Dashed lines show predictions for the same sequences obtained using a null model trained on pluripotent chromatin accessibility. **b)** Scatter plots comparing ChromBPNet-predicted log₂ accessibility (x-axis) to MPRA-measured activity (y-axis) for the enhancement (left) and ablation (right) trajectories. Points correspond to intermediate sequences along each trajectory; the starting sequence (black circle), best-performing intermediate for the corresponding objective (star), and final intermediate (red or blue circle) are indicated, and the color gradient denotes the number of mutations introduced. Horizontal dotted lines (with shading) denote the activity of the endogenous mouse CRE (green) and the minimal-promoter (minP) background control (blue). Vertical green dotted lines denote the predicted accessibility of the endogenous mouse CRE. Spearman correlation coefficients (ρ) between predicted accessibility and MPRA activity are shown for each combination of CRE and training objective.

**Supplementary Figure 11.**
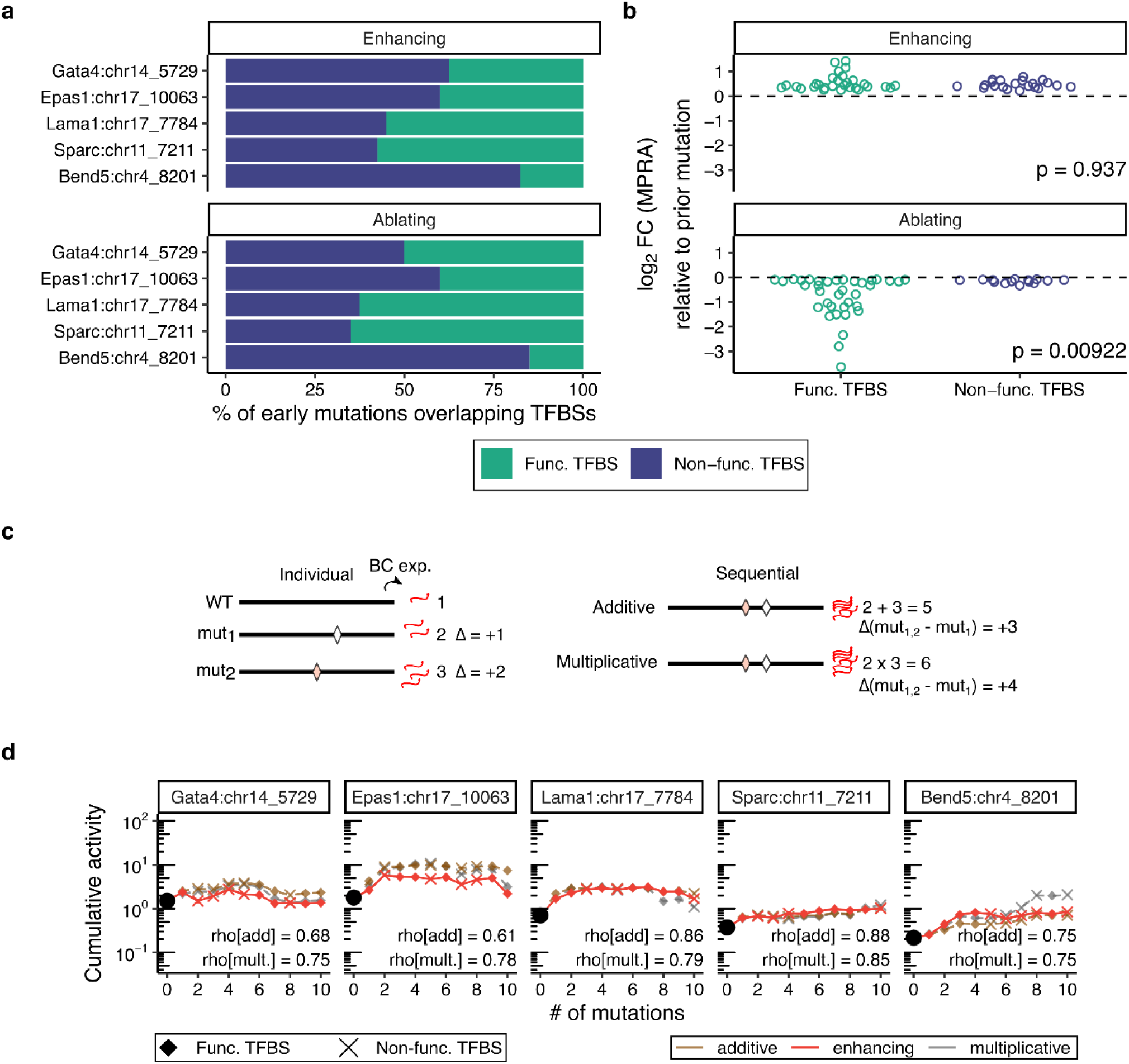
Preferential targeting of functional TFBSs and epistasis during model-guided modulation of parietal endoderm CREs. **a)** Composition of the first ten model-nominated mutations for each CRE under the enhancement and ablation objectives, stratified by whether mutations overlap functional TFBSs (green) or non-TFBS positions (purple) in the extant mouse sequence. **b)** Log₂ fold-changes in MPRA activity contributed by each of the first ten mutations, measured relative to the immediately preceding intermediate sequence, shown separately for enhancement and ablation objectives and stratified by mutations overlapping functional TFBSs (green) or non-TFBS positions (purple). P-values indicate Wilcoxon rank-sum tests comparing activity changes between TFBS-overlapping and non-TFBS mutations. **c)** Schematic of the additive and multiplicative models used to estimate the combined effects of successive mutations. In the additive model, mutational effects on reporter output are assumed to combine additively, whereas in the multiplicative model they are assumed to combine multiplicatively. **d)** Cumulative MPRA activity for the first ten mutations for each of the five CREs, compared to expectations from the additive (brown dashed) and multiplicative (grey dashed) models. MPRA-measured activity is shown as solid red lines. Error bars denote the standard deviation across three biological replicates. The black circle marks the activity of the endogenous mouse CRE. Symbols indicate whether individual mutations overlap functional TFBSs (diamonds) or non-TFBS positions (crosses). Spearman correlation coefficients (ρ) between observed and expected activities are shown for each CRE and model.

## SUPPLEMENTARY NOTE

**Supplementary Note 1. Comparison of four sequence-based modeling strategies.** To assess how well enhancer activity across the mammalian phylogeny can be predicted directly from sequence, we evaluated four sequence-based modeling strategies of increasing complexity across five parietal endoderm CREs, including: **(i)** a simple “functional motif count”, obtained by projecting TFBSs experimentally mapped in extant mouse onto orthologs; **(ii)** a generalized linear model (GLM) trained on predicted TF binding affinities; **(iii)** a gapped k-mer support vector machine model, gkm-SVM^45^, trained on parietal endoderm accessibility; and **(iv)** a deep convolutional neural network (CNN), ChromBPNet^46^, trained on the same accessibility profiles. Evaluating these models in parallel allowed us to assess how much of the striking functional diversity observed across orthologs of each CRE can be explained by progressively richer representations of sequence information.

### Functional motif counts

As a baseline approach, we projected mouse TFBSs mapped by saturation mutagenesis^38^ onto all orthologs (7-15 mouse TFBSs per CRE, and counted those retaining comparable or greater predicted affinity. Most orthologs contained fewer sites than the mouse reference, and the number of sites was moderately correlated with MPRA activity (ρ = 0.15-0.78). However, the impact of site loss differed by element: *Gata4* CRE orthologs generally required near-complete retention of equivalent TFBSs to maintain activity, while *Epas1* and *Lama1* CRE orthologs often preserved or even exceeded mouse activity despite losing multiple mouse motifs (**Extended Data Fig. 1a**) These results indicate that simple motif composition partly explains cross-species functional variation, and suggests substantial heterogeneity in the rigidity versus flexibility across elements.

### GLMs based on predicted TF binding affinity

To incorporate quantitative motif strength, we fit a GLM for each CRE using summed predicted affinities for key parietal endoderm TFs as predictors. Relative to simple counts, these models roughly doubled correlation with MPRA activity for 3 of the 5 CREs (ρ = 0.30-0.61; **Extended Data Fig. 1b**), indicating that variation in TFBS affinity also contributes meaningfully to functional divergence across orthologs. Consistent with saturation mutagenesis data, AP-1 affinity most strongly predicted activity for *Epas1* CRE orthologs, while *Sox17* affinity best explained variation among *Gata4* CRE orthologs (**Extended Data Fig. 1c**).

### gkm-SVM trained on chromatin accessibility

As a third approach, we trained a model using gkm-SVM^45^, a support vector machine framework that uses gapped k-mers, on pseudobulk parietal endoderm ATAC-seq peaks from our previous scATAC-seq dataset^34^. Unlike the motif-based models above, gkm-SVM is trained to predict chromatin accessibility, rather than enhancer activity, directly from sequence. When applied to evolutionary and ancestral CRE orthologs, gkm-SVM predictions showed weak to moderate correlations with MPRA activity in four of five cases (ρ = 0.01–0.57; **Extended Data Fig. 1d**). Notably, gkm-SVM outperformed motif count and GLM approaches for the *Lama1* (ρ = 0.57) and *Bend5* (ρ = 0.41) CREs, indicating that k-mer–based sequence features capture aspects of regulatory variation not represented by motif-centric models.

### ChromBPNet modeling of chromatin accessibility

To leverage recent advances in deep learning-based modeling of regulatory grammar, we applied ChromBPNet^46^, a CNN for modeling chromatin accessibility, trained on the same pseudobulked parietal endoderm scATAC-seq data as the gkm-SVM model parietal endoderm, as well as data from other mEB-derived lineages (**Extended Data Fig. 2a**). Predicted and observed accessibility were well-correlated for each pluripotent and germ layer lineages, both on consensus peaks (ρ = 0.60-0.83) and differential accessible peaks (ρ = 0.61-0.78) (**Extended Data Fig. 3a-b**). Moreover, ChromBPNet predicted accessibility well over genomic regions corresponding to 4 of the 5 CREs tested (all but the *Sparc* CRE; **Extended Data Fig. 3c**). Despite being a predictor of accessibility, ChromBPNet performed surprisingly well when as a predictor of MPRA activity (average of accessibility profile across embedding in 100 random sequences), (ρ = 0.35-0.64; **Extended Data Fig. 3d**).

### Comparative performance of sequence models

While all four approaches recovered meaningful sequence–function relationships, ChromBPNet provided the most consistent and generalizable performance across CREs on our task. Motif counts provide an intuitive baseline but are overly reductionist, collapsing each ortholog to the presence or absence of a small set of mouse-mapped TFBSs, missing syntax, compensatory changes, and gains of novel motifs. GLMs improved predictions for several CREs, but are trained “per CRE” and thus do not generalize to new elements and are further limited by linear assumptions. gkm-SVM, trained once on genome-wide accessibility, avoids CRE-specific overfitting but exhibits variable performance across elements. In contrast, ChromBPNet achieved moderately strong correlations across all five CREs, offering the most stable and generalizable sequence-to-activity predictions. This, coupled with its compatibility with nucleotide-resolution interpretation tools, motivated us to use it as our primary model for downstream analyses.

### Base-resolution interpretation and motif perturbation analyses

We also leveraged ChromBPNet’s base-resolution interpretations to query what sequence features drive accessibility across species. Comparing mouse and rat *Gata4* CREs, the model highlighted the loss of the *Sox17* motif in rat (**Extended Data Fig. 2c**), a site gained along the mouse lineage after divergence from Anc38 (**Fig. 2f-g**; **Supplementary Fig. 5a**) and coinciding with gain of function as measured by MPRA. Similarly, model interpretations for the mouse and African wild dog *Epas1* CREs emphasized the conserved Foxa2-Sox17-Gata4/6 triplet as the dominant contributor to predicted accessibility in both species (**Extended Data Fig. 2d**). ChromBPNet also flagged specific substitutions within this module that matched effects measured by saturation mutagenesis in mouse, including their impacts on predicted TFBS affinity (**Extended Data Fig. 2e**). In the African wild dog ortholog, these substitutions decreased predicted Foxa2 and Gata4/6 affinity (3.6-fold and 3.7-fold, respectively) while increasing Sox17 affinity 2.3-fold, potentially examples of buffering via compensatory changes.

Finally, we performed *in silico* insertions of key parietal endoderm TF motifs into non-accessible background sequences, and evaluated their impact on model-predicted chromatin accessibility. Particularly for Fox, Sox and Gata motifs, *in silico* insertion yielded “peak gains” with the parietal endoderm model but not other germ layer models (**Extended Data Fig. 4a**). Furthermore, analysis of learned motifs using TF-MoDISco^83^ revealed high concordance between models trained on pseudobulk parietal endoderm scATAC-seq and PYS-2 bulk ATAC-seq data (**Extended Data Fig. 4b**).

These complementary evaluators of sequence syntax highlight how diverse regulatory architectures can support transcriptional activity in PYS-2 cells. Elements such as the *Gata4* CRE appear to rely on a precise grammar structure that constrains sequence evolution, whereas others, like *Epas1*, reach activity once a threshold of overall binding affinity is reached—a pattern consistent with a billboard-like regulatory model^84^. Although chromatin accessibility alone does not determine enhancer activity, it remains a prerequisite for regulatory potential^85^, providing a useful surrogate for modeling sequence features associated with activation. Accordingly, we used the accessibility-trained model to identify and quantify base-resolution contributions to transcriptional activity in the analyses that follow.

## EXTENDED DATA FIGURES 1-4

**Extended Data Figure 1.**
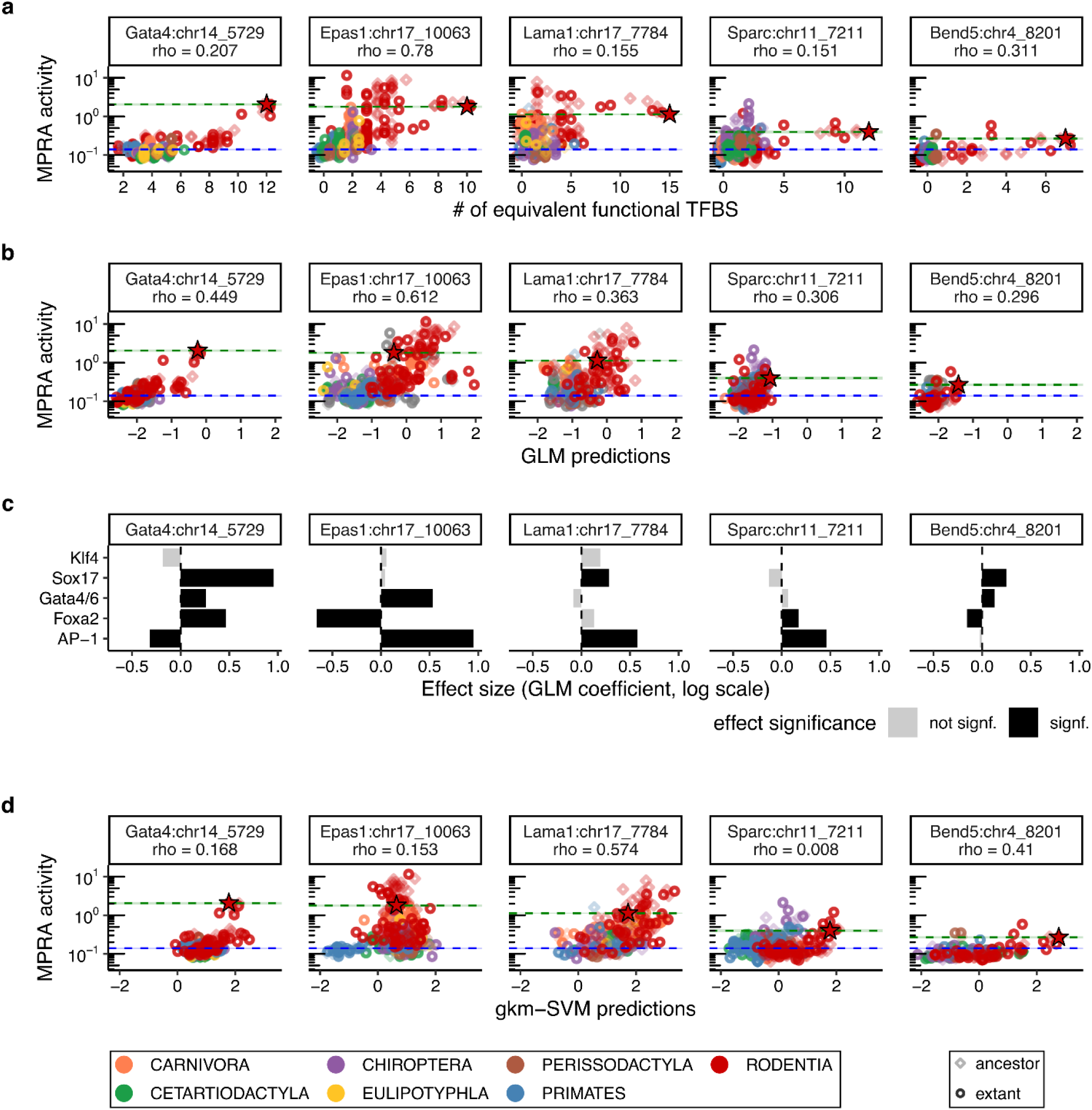
Sequence-based predictors of CRE activity. a-b) Scatterplots for each model CRE of MPRA activity (y-axes) vs. the number of mouse TFBSs with predicted affinities greater than or equal to the endogenous mouse sites (TFBS affinity > 0.05) **(a)** or generalized linear model (GLM) predictions **(b)** (x-axes). Spearman’s ρ values report correlation. Dotted lines indicate endogenous mouse CRE (green) and minP background (blue) activity levels, respectively. Diamonds correspond to ancestral orthologs, circles to extant orthologs, and red stars to the endogenous mouse CRE. Colors correspond to phylogenetic orders (key at bottom of figure). **c)** Effect sizes for summed predicted TFBS affinities (TFBS affinity > 0.05) estimated by generalized linear models (GLMs) for each key TF motif in each CRE. Gray and black bars denote non-significance and significance, respectively (p-value < 0.05 using the Wald test). **d)** Scatterplots for each model CRE of MPRA activity (y-axes) vs. parietal-endoderm-trainedgkm-SVM predictions (x-axes). Presentation details match panels **a-b**.

**Extended Data Figure 2.**
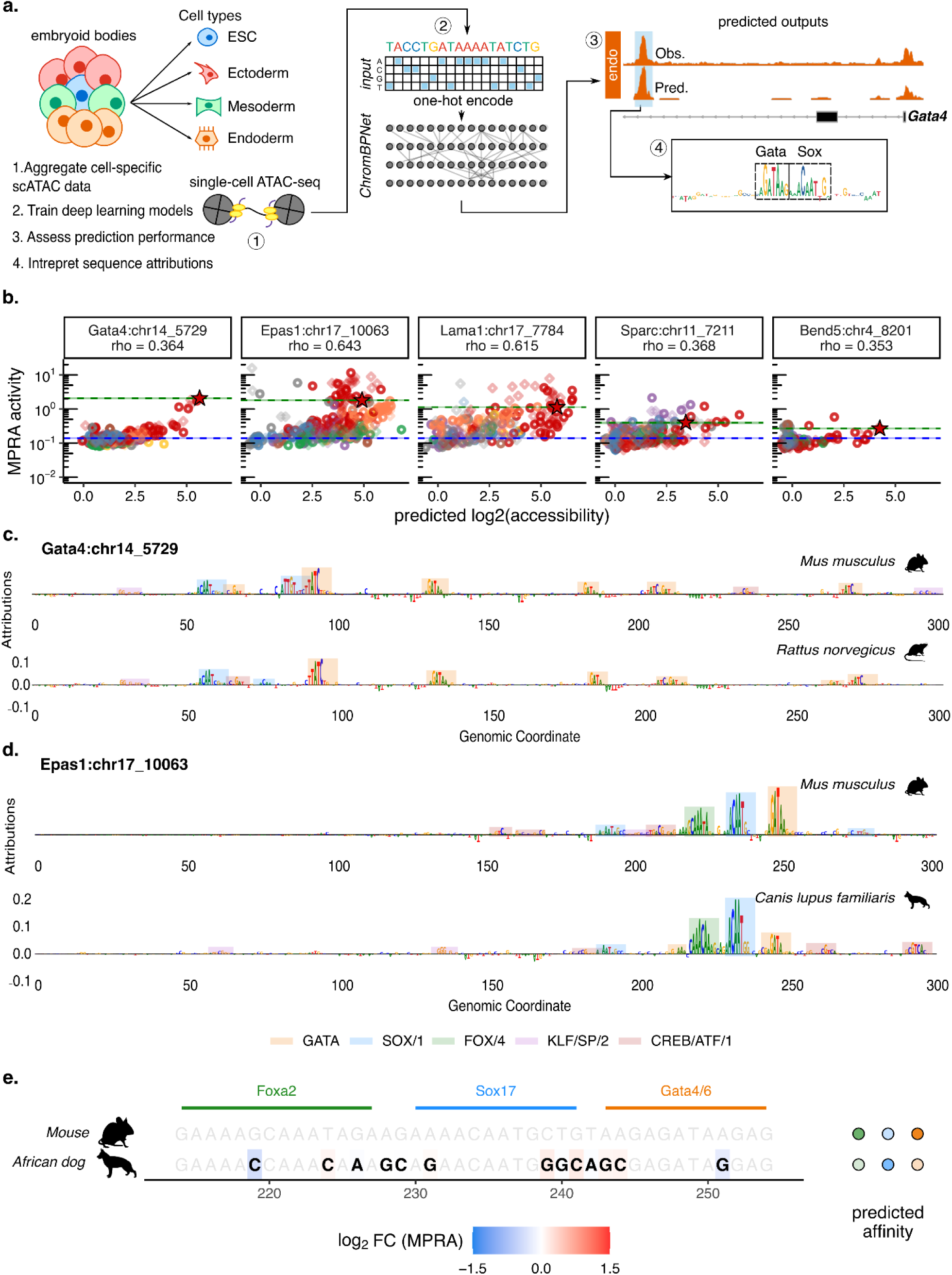
ChromBPNet-based interpretation of parietal endoderm CRE orthologs. **a)** Schematic overview of ChromBPNet model training, evaluation, and sequence interpretation from mouse parietal endoderm scATAC-seq data. **b)** Scatterplots for each model CRE of MPRA activity (y-axes) vs. ChromBPNet predictions. Spearman’s ρ values report correlation. Dotted lines indicate endogenous mouse CRE (green) and minP background (blue) activity levels, respectively. Diamonds correspond to ancestral orthologs, circles to extant orthologs, and red stars to the endogenous mouse CRE. Colors correspond to phylogenetic orders (key at bottom of **Extended Data Fig. 1**). **c-d)** ChromBPNet profile contribution scores for mouse Gata4:chr14_5729 CRE and its rat ortholog **(c)** and mouse *Epas1*:chr17_10063 CRE and its African dog ortholog **(d)**. Color-shaded boxes below correspond to mapped functional TFBS. **e)** Left: Nucleotide-resolution multiple sequence alignment (MSA) focused on the conserved Foxa2-Sox17-Gata4/6 module within the mouse Epas1:chr17_10063 CRE and the corresponding region in the African wild dog ortholog. Bolded nucleotides in black text denote substitutions relative to the mouse *Epas1* CRE, colored by log₂ fold-change in saturation-mutagenesis MPRA activity, while wild-type nucleotides are unbolded (grey text). Right: Predicted TFBS affinities for each ortholog, color-coded by relative affinity.

**Extended Data Figure 3.**
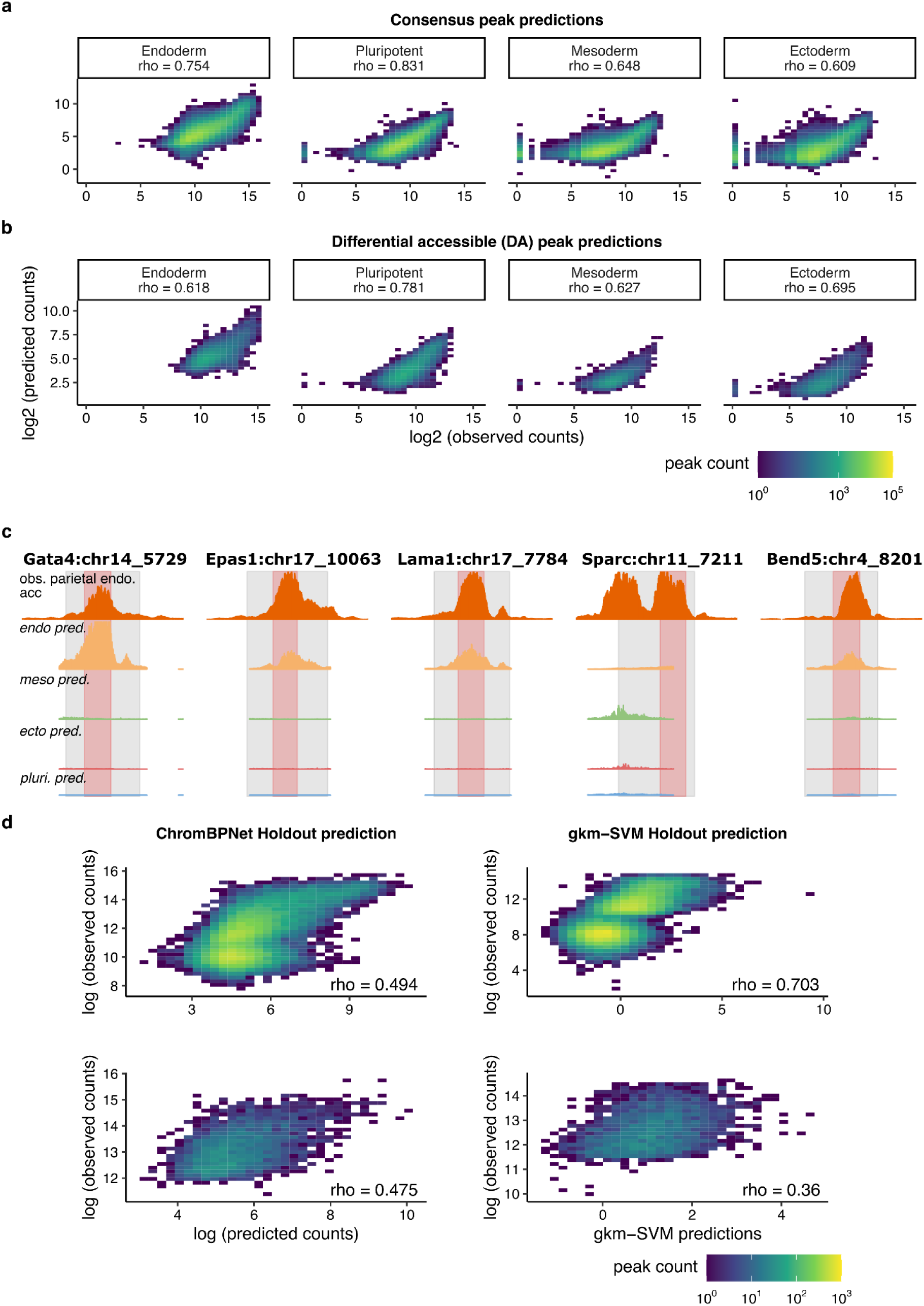
ChromBPNet modeling of chromatin accessibility. a-b) Scatterplots showing log-scaled observed (x-axes) vs. log-scaled ChromBPNet predicted (y-axes) read counts across mEB consensus peaks (*n* = 365,392) **(a)** or cell-specific differentially accessible (DA) consensus peaks (bottom; *n* = 56,795) **(b)** for each trained cell type model. Spearman’s ρ values report correlation. **c)** Pseudo-bulk read pileup for parietal endoderm scATAC observed data (top row) vs. ChromBPNet predicted profiles for each cell type (bottom rows) for each model CRE (columns). Shaded gray regions indicate the genomic coordinates of the original full-length CREs as validated by scQer^34^ (520 bp to 1.7 kb in length). Shaded red regions correspond to coordinates of 300-bp maximum activity tile used for downstream experiments. **d)** Scatterplots showing log-scaled observed (x-axes) vs. log-scaled ChromBPNet (left column) or gkm-SVM (right column) predicted (y-axes) read counts across all held-out chromosomal test peaks (top row; *n* = 30,540) and parietal endoderm-specific differentially accessible (DA) peaks (bottom row; n = 3,099 and 2,904 for ChromBPNet and gkm-SVM, respectively). Both models were trained on the same parietal endoderm pseudo-bulk scATAC-seq data. Spearman’s ρ values report correlation.

**Extended Data Figure 4.**
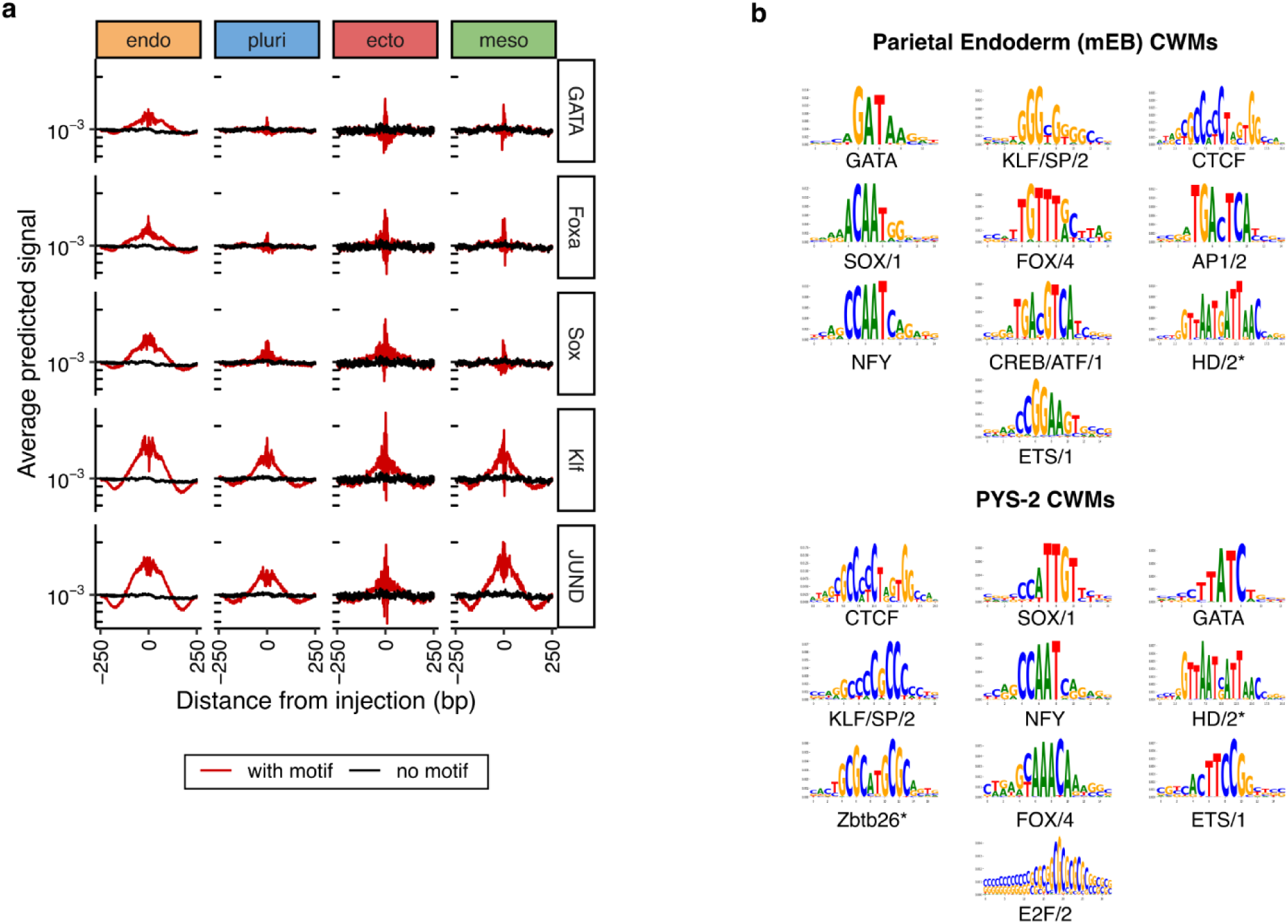
Model-learned motifs from mouse parietal endoderm ATAC data. **a)** Marginal footprints of inserting parietal endoderm-specific TF motifs (columns) into genomic background sequences, using ChromBPNet predicted profiles across all mEB cell type models (rows). Red and black indicate motif insertion and no insertion, respectively. **b)** Top 10 TF-MODISCO seqlets (ranked by the number of predictive motif instances and named by motif families) derived from contribution weighted matrices (CWMs) from ChromBPNet models trained on pseudo-bulk scATAC data from parietal endoderm in mEBs (top) and bulk ATAC data from PYS-2 cell lines (bottom). Asterisks indicate motifs identified by TOMTOM that failed the statistical threshold (*i.e.* q > 0.05).

## REFERENCES

1. Christmas, M. J. et al. Evolutionary constraint and innovation across hundreds of placental mammals. Science 380, eabn3943 (2023).

2. Armstrong, J. et al. Progressive Cactus is a multiple-genome aligner for the thousand-genome era. Nature 587, 246–251 (2020).

3. Carroll, S. B. Evo-devo and an expanding evolutionary synthesis: a genetic theory of morphological evolution. Cell 134, 25–36 (2008).

4. Nord, A. S. et al. Rapid and pervasive changes in genome-wide enhancer usage during mammalian development. Cell 155, 1521–1531 (2013).

5. Stergachis, A. B. et al. Conservation of trans-acting circuitry during mammalian regulatory evolution. Nature 515, 365–370 (2014).

6. Wilson, M. D. et al. Species-specific transcription in mice carrying human chromosome 21. Science 322, 434–438 (2008).

7. Villar, D. et al. Enhancer evolution across 20 mammalian species. Cell 160, 554–566 (2015).

8. Kasowski, M. et al. Variation in transcription factor binding among humans. Science 328, 232–235 (2010).

9. King, M. C. & Wilson, A. C. Evolution at two levels in humans and chimpanzees: Their macromolecules are so alike that regulatory mutations may account for their biological differences. Science 188, 107–116 (1975).

10. Kvon, E. Z. et al. Comprehensive in vivo interrogation reveals phenotypic impact of human enhancer variants. Cell 180, 1262–1271.e15 (2020).

11. Kircher, M. et al. A general framework for estimating the relative pathogenicity of human genetic variants. Nat. Genet. 46, 310–315 (2014).

12. Jindal, G. A. & Farley, E. K. Enhancer grammar in development, evolution, and disease: dependencies and interplay. Dev. Cell 56, 575–587 (2021).

13. Kvon, E. Z. et al. Progressive loss of function in a limb enhancer during snake evolution. Cell 167, 633–642.e11 (2016).

14. Hare, E. E., Peterson, B. K., Iyer, V. N., Meier, R. & Eisen, M. B. Sepsid even-skipped enhancers are functionally conserved in Drosophila despite lack of sequence conservation. PLoS Genet. 4, e1000106 (2008).

15. Wong, E. S. et al. Deep conservation of the enhancer regulatory code in animals. Science 370, eaax8137 (2020).

16. McDonald, J. M. C. & Reed, R. D. Beyond modular enhancers: new questions in cis-regulatory evolution. Trends Ecol. Evol. 39, 1035–1046 (2024).

17. Zoonomia Consortium. A comparative genomics multitool for scientific discovery and conservation. Nature 587, 240–245 (2020).

18. Pollard, K. S., Hubisz, M. J., Rosenbloom, K. R. & Siepel, A. Detection of nonneutral substitution rates on mammalian phylogenies. Genome Res. 20, 110–121 (2010).

19. ENCODE Project Consortium. An integrated encyclopedia of DNA elements in the human genome. Nature 489, 57–74 (2012).

20. Kosicki, M. et al. VISTA Enhancer browser: an updated database of tissue-specific developmental enhancers. Nucleic Acids Res. 53, D324–D330 (2025).

21. Berthelot, C., Villar, D., Horvath, J. E., Odom, D. T. & Flicek, P. Complexity and conservation of regulatory landscapes underlie evolutionary resilience of mammalian gene expression. *Nat*. Ecol. Evol. 2, 152–163 (2018).

22. Patwardhan, R. P. et al. Massively parallel functional dissection of mammalian enhancers in vivo. Nat. Biotechnol. 30, 265–270 (2012).

23. Pennacchio, L. A. et al. In vivo enhancer analysis of human conserved non-coding sequences. Nature 444, 499–502 (2006).

24. Uebbing, S. et al. Massively parallel discovery of human-specific substitutions that alter enhancer activity. Proc Natl Acad Sci U S A 118, (2021).

25. Barbadilla-Martínez, L., Klaassen, N., van Steensel, B. & de Ridder, J. Predicting gene expression from DNA sequence using deep learning models. Nat. Rev. Genet. 1–15 (2025).

26. Kim, S. & Wysocka, J. Deciphering the multi-scale, quantitative cis-regulatory code. Mol. Cell 83, 373–392 (2023).

27. Minnoye, L. et al. Cross-species analysis of enhancer logic using deep learning. Genome Res. 30, 1815–1834 (2020).

28. Oh, J. W. & Beer, M. A. Gapped-kmer sequence modeling robustly identifies regulatory vocabularies and distal enhancers conserved between evolutionarily distant mammals. Nat. Commun. 15, 6464 (2024).

29. Hochberg, G. K. A. & Thornton, J. W. Reconstructing ancient proteins to understand the causes of structure and function. Annu. Rev. Biophys. 46, 247–269 (2017).

30. Starr, T. N., Picton, L. K. & Thornton, J. W. Alternative evolutionary histories in the sequence space of an ancient protein. Nature 549, 409–413 (2017).

31. Klein, J. C., Keith, A., Agarwal, V., Durham, T. & Shendure, J. Functional characterization of enhancer evolution in the primate lineage. Genome Biol. 19, 99 (2018).

32. Hecker, N. et al. Enhancer-driven cell type comparison reveals similarities between the mammalian and bird pallium. Science 387, eadp3957 (2025).

33. Inoue, F. Massively parallel reporter assays: from barcodes to biology. Nat Rev Genet (2026) doi:10.1038/s41576-026-00944-4.

34. Lalanne, J.-B. et al. Multiplex profiling of developmental cis-regulatory elements with quantitative single-cell expression reporters. Nat. Methods 21, 983–993 (2024).

35. Lehman, J. M., Speers, W. C., Swartzendruber, D. E. & Pierce, G. B. Neoplastic differentiation: characteristics of cell lines derived from a murine teratocarcinoma. J. Cell. Physiol. 84, 13–27 (1974).

36. Brown, K. et al. A comparative analysis of extra-embryonic endoderm cell lines. PLoS One 5, e12016 (2010).

37. Argelaguet, R. et al. Decoding gene regulation in the mouse embryo using single-cell multi-omics. bioRxiv 2022.06.15.496239 (2022) doi:10.1101/2022.06.15.496239.

38. Lalanne, J.-B. et al. Multi-scale dissection, compaction, and derivatization of mammalian developmental enhancers. bioRxiv.

39. Weberling, A. et al. Primitive to visceral endoderm maturation is essential for mouse epiblast survival beyond implantation. iScience 28, 111671 (2025).

40. Stary, M. et al. Parietal endoderm secreted SPARC promotes early cardiomyogenesis in vitro. Exp. Cell Res. 310, 331–343 (2005).

41. Mouse Genome Sequencing Consortium et al. Initial sequencing and comparative analysis of the mouse genome. Nature 420, 520–562 (2002).

42. Kircher, M. et al. Saturation mutagenesis of twenty disease-associated regulatory elements at single base-pair resolution. Nat. Commun. 10, 3583 (2019).

43. IGVF Consortium. Deciphering the impact of genomic variation on function. Nature 633, 47–57 (2024).

44. Moore, J. E. et al. An expanded Registry of candidate cis-Regulatory Elements for studying transcriptional regulation. bioRxivorg 2024.12.26.629296 (2024) doi:10.1101/2024.12.26.629296.

45. Ghandi, M., Lee, D., Mohammad-Noori, M. & Beer, M. A. Enhanced regulatory sequence prediction using gapped k-mer features. PLoS Comput. Biol. 10, e1003711 (2014).

46. Pampari, A. et al. ChromBPNet: bias factorized, base-resolution deep learning models of chromatin accessibility reveal cis-regulatory sequence syntax, transcription factor footprints and regulatory variants. bioRxivorg 2024.12.25.630221 (2025) doi:10.1101/2024.12.25.630221.

47. Inoue, F. & Ahituv, N. Decoding enhancers using massively parallel reporter assays. Genomics 106, 159–164 (2015).

48. Taskiran, I. I. et al. Cell-type-directed design of synthetic enhancers. Nature 626, 212–220 (2024).

49. Martyn, G. E. et al. Rewriting regulatory DNA to dissect and reprogram gene expression. Cell (2025) doi:10.1016/j.cell.2025.03.034.

50. de Boer, C. G. & Taipale, J. Hold out the genome: a roadmap to solving the cis-regulatory code. Nature 625, 41–50 (2024).

51. Jindal, G. A. et al. Single-nucleotide variants within heart enhancers increase binding affinity and disrupt heart development. Dev. Cell 58, 2206–2216.e5 (2023).

52. Romano, L. A. & Wray, G. A. Conservation of Endo16 expression in sea urchins despite evolutionary divergence in both cis and trans-acting components of transcriptional regulation. Development 130, 4187–4199 (2003).

53. Swanson, C. I., Schwimmer, D. B. & Barolo, S. Rapid evolutionary rewiring of a structurally constrained eye enhancer. Curr. Biol. 21, 1186–1196 (2011).

54. Ruvinsky, I. & Ruvkun, G. Functional tests of enhancer conservation between distantly related species. Development 130, 5133–5142 (2003).

55. Wratten, N. S., McGregor, A. P., Shaw, P. J. & Dover, G. A. Evolutionary and functional analysis of the *tailless* enhancer in *Musca domestica* and *Drosophila melanogaster*. Evol. Dev. 8, 6–15 (2006).

56. Moses, A. M. et al. Large-scale turnover of functional transcription factor binding sites in Drosophila. PLoS Comput. Biol. 2, e130 (2006).

57. Bradley, R. K. et al. Binding site turnover produces pervasive quantitative changes in transcription factor binding between closely related Drosophila species. PLoS Biol. 8, e1000343 (2010).

58. Qiu, C. et al. Evolutionary transfer learning enables organism-wide inference of mammalian enhancer landscapes. bioRxiv 2026.04.07.717039 (2026) doi:10.64898/2026.04.07.717039.

59. Chen, A., Chen, D. & Chen, Y. Advances of DNase-seq for mapping active gene regulatory elements across the genome in animals. Gene 667, 83–94 (2018).

60. Ernst, J. et al. Genome-scale high-resolution mapping of activating and repressive nucleotides in regulatory regions. Nat. Biotechnol. 34, 1180–1190 (2016).

61. Agarwal, V. et al. Massively parallel characterization of transcriptional regulatory elements. Nature 639, 411–420 (2025).

62. Gosai, S. J. et al. Machine-guided design of cell-type-targeting cis-regulatory elements. Nature 634, 1211–1220 (2024).

63. Murphy, A., Durán, A. & Koo, P. Adapting AlphaGenome to MPRA data. Genomics x AI https://genomicsxai.github.io/blogs/2026-002/ (2026).

64. Pinglay, S. et al. Mammalian genome writing: Unlocking new length scales for genome engineering. Cell 189, 356–374 (2026).

65. Hoose, A., Vellacott, R., Storch, M., Freemont, P. S. & Ryadnov, M. G. DNA synthesis technologies to close the gene writing gap. Nat. Rev. Chem. 7, 144–161 (2023).

66. Koo, P. K. Decoding the regulatory genome with large-scale deep learning. Nat. Rev. Genet. 27, 117 (2026).

67. Blanchette, M., Green, E. D., Miller, W. & Haussler, D. Reconstructing large regions of an ancestral mammalian genome in silico. Genome Res 14, 2412–2423 (2004).

68. Domcke, S. et al. A human cell atlas of fetal chromatin accessibility. Science 370, eaba7612 (2020).

69. Corces, M. R. et al. An improved ATAC-seq protocol reduces background and enables interrogation of frozen tissues. Nat. Methods 14, 959–962 (2017).

70. Quinlan, A. R. & Hall, I. M. BEDTools: a flexible suite of utilities for comparing genomic features. Bioinformatics 26, 841–842 (2010).

71. Kent: UCSC Genome Browser Source Tree. Stable Branch: ‘Beta’. (Github).

72. Li, H. et al. The Sequence Alignment/Map format and SAMtools. Bioinformatics 25, 2078–2079 (2009).

73. Feng, J., Liu, T., Qin, B., Zhang, Y. & Liu, X. S. Identifying ChIP-seq enrichment using MACS. Nat. Protoc. 7, 1728–1740 (2012).

74. Sequence_align: Efficient Implementations of Needleman-Wunsch and Other Sequence Alignment Algorithms Written in Rust with Python Bindings via PyO3. (Github).

75. Krueger, F. TrimGalore: A Wrapper around Cutadapt and FastQC to Consistently Apply Adapter and Quality Trimming to FastQ Files, with Extra Functionality for RRBS Data. (Github).

76. Zhang, J., Kobert, K., Flouri, T. & Stamatakis, A. PEAR: a fast and accurate Illumina Paired-End reAd mergeR. Bioinformatics 30, 614–620 (2014).

77. Langmead, B. & Salzberg, S. L. Fast gapped-read alignment with Bowtie 2. Nat. Methods 9, 357–359 (2012).

78. Keukeleire, P. et al. Using individual barcodes to increase quantification power of massively parallel reporter assays. BMC Bioinformatics 26, 52 (2025).

79. Rube, H. T. et al. Prediction of protein-ligand binding affinity from sequencing data with interpretable machine learning. Nat. Biotechnol. 40, 1520–1527 (2022).

80. Katoh, K., Misawa, K., Kuma, K.-I. & Miyata, T. MAFFT: a novel method for rapid multiple sequence alignment based on fast Fourier transform. Nucleic Acids Res. 30, 3059–3066 (2002).

81. Xu, S., et al. Ggtree: A serialized data object for visualization of a phylogenetic tree and annotation data. Imeta 1, e56 (2022).

82. Chung, J. H., Bell, A. C. & Felsenfeld, G. Characterization of the chicken beta-globin insulator. Proc. Natl. Acad. Sci. U. S. A. 94, 575–580 (1997).

83. Shrikumar, A. et al. Technical note on Transcription Factor Motif Discovery from Importance Scores (TF-MoDISco) version 0.5.6.5. arXiv [cs.LG] (2018).

84. Kulkarni, M. M. & Arnosti, D. N. Information display by transcriptional enhancers. Development 130, 6569–6575 (2003).

85. Thurman, R. E. et al. The accessible chromatin landscape of the human genome. Nature 489, 75–82 (2012).

